# The retroelement-derived human protein PEG10 is a regulator of mRNA splicing in neurons

**DOI:** 10.64898/2026.05.21.727000

**Authors:** A.M. Matthews, A.M. Whiteley

## Abstract

Retroelements, including retrotransposons, endogenous retroviruses, and their fragments, as well as rare co-opted or domesticated retroelements, can contribute to neurodegenerative disorders and aging through modulation of gene expression and induction of neuroinflammation. Paternally Expressed Gene 10 (PEG10) is a retroelement-derived human gene that has recently been identified as a putative driver of Amyotrophic Lateral Sclerosis (ALS) and Angelman’s Syndrome. PEG10 has been reported to bind nucleic acid and undergoes a complex self-processing pathway that results in gene expression changes when the protein accumulates in cells. Here, we report that PEG10 has selectivity for binding U/G-rich RNAs and influences widespread gene expression changes. PEG10 overexpression mimics the loss of TDP-43 in broad changes to gene expression, including dysregulation of mRNA splicing pathways. Specific changes to mRNA splicing were largely unique between TDP-43 knockdown and PEG10 overexpression, as classic TDP-43 targets including *STMN2* were not altered by PEG10. Instead, we identified a unique role for PEG10 in regulating splicing of *neuregulin 3 (NRG3)*, a ligand for the neuronal receptor ERBB4. In SH-SY5Y cells and in human neurons overexpressing PEG10, NRG3 protein levels were decreased along cellular processes, suggesting that these cells are less competent at signaling through the NRG3/ERBB4 axis. Using human patient data, we observed similar changes to *NRG3* splicing in *UBQLN2*-mediated ALS, where PEG10 is accumulated, as well as in some cases of sporadic ALS. In conclusion, the retroelement-derived gene PEG10 plays an unexpected role in regulating splicing of neuronal transcripts, which mimics some of the transcript changes observed in human ALS patient samples. Ultimately, this work has implications for the study of PEG10, and mRNA splicing in neurological diseases associated with elevated PEG10 abundance.

**Highlights:**

- PEG10 NC expression influences abundance of transcripts implicated in ALS
- PEG10 NC expression leads to an exon skipping event in *neuregulin 3 (NRG3)*
- NRG3 expression is decreased along dendrites of PEG10 NC expressing human neurons
- Expression of PEG10 NC mimics changes observed in human ALS

**Graphical Abstract:** 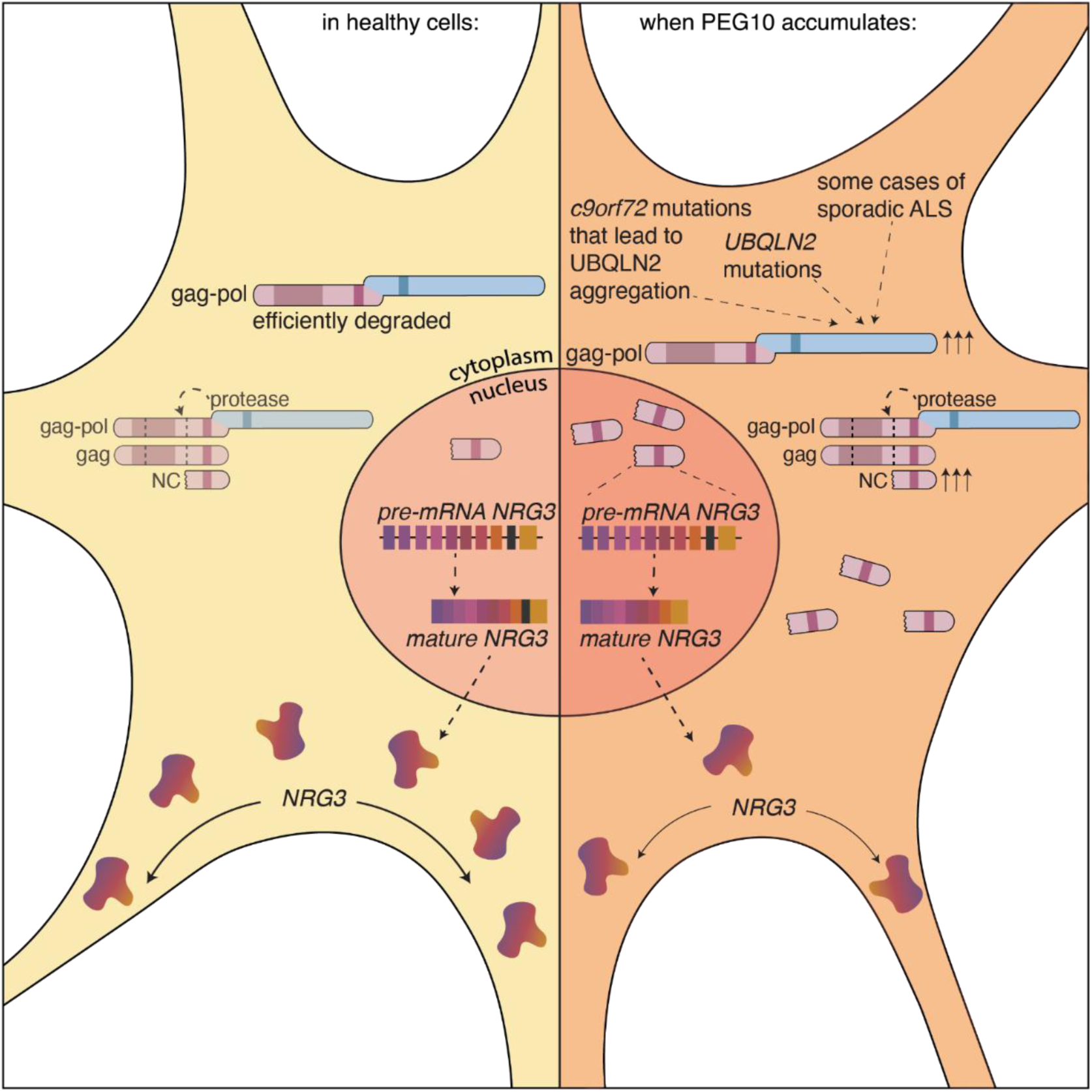

## Introduction

Retroelements are frequently derepressed in aging and neurological disease, and can contribute to the disease process through increased inflammatory signaling, genome instability, and induction of antiviral responses^1–4^. Amyotrophic Lateral Sclerosis (ALS) is a fatal neurodegenerative disease characterized by a progressive loss of motor function due to dysfunction and death of upper and lower motor neurons^5,6^. In some cases, ALS is complicated by the development of frontotemporal dementia (FTD), resulting in a spectrum of symptoms from exclusive motor neuron degeneration to a disease course dominated by cortical degeneration^7,8^. 10% of ALS cases are familial (fALS) and are linked to genetic mutation of over three dozen genes^9^, while 90% of cases are sporadic (sALS) and occur with no clear cause or family history^5^. Despite the complex origins of disease, ALS cases are tied together by hallmarks of neuronal dysfunction, including failed protein degradation, cytoskeletal abnormalities, and dysfunction of RNA binding proteins, including the critical splice regulator TDP-43^6,10–14^. Upregulation of retroelement gene expression is also observed in a subset of ALS patients, and correlates with the level of dysfunction of TDP-43^15–17^. TDP-43 silences expression of transposable elements (TEs), suggesting that TE derepression and its downstream effects are a result of TDP-43 dysfunction in disease^18–21^. However, there is also evidence that TE expression can influence TDP-43 function, thereby indirectly contributing to splicing defects and other molecular hallmarks of ALS^20^.

Previous work from mouse models and human patient tissues showed that a domesticated retroelement, Paternally Expressed Gene 10 (PEG10), is dramatically accumulated in *UBQLN2*-mediated familial ALS-FTD^22^, as well as sporadic ALS^23^, and Angelman’s Syndrome^24^. In the case of ALS, it is thought that mutation of *UBQLN2* alleles in fALS^25^ lead to failed PEG10 degradation^22,23,26,27^. However, aggregation of UBQLN2 is also observed in *c9orf72*-mediated ALS^28^, *FUS*-mediated ALS^29^, and in some cases of sporadic ALS^28,30,31^, likely leading to PEG10 accumulation via dysfunctional degradation by UBQLN2.

Despite its domestication in placental mammals over 200 million years ago^32–36^, PEG10 retains a number of retroelement-like features that implicate it as a potential driver of cellular dysfunction. Like retroviruses and retrotransposons, PEG10 utilizes programmed ribosomal frameshifting to generate distinct gag and gag-pol proteins from the same mRNA^37,38^. The pol region of PEG10 retains an active retroviral protease domain that is capable of cleaving both the gag-pol and gag proteins (Figure 1A)^23,39^. Proteolytic cleavage of the abundant gag protein liberates a 68 amino acid nucleic acid-binding fragment which then localizes to the nucleus^23^. This unique regulatory pathway suggests that accumulation of the protease-containing gag-pol protein may reveal new roles of PEG10 in the regulation of nucleic acids, especially in the context of neurological disease. Here, we describe a novel role for PEG10 in regulating mRNA splicing and contributing to dysregulation of neuronal signaling pathways, which has implications for when PEG10 is accumulated in neurological diseases, including ALS.

**Figure 1.**
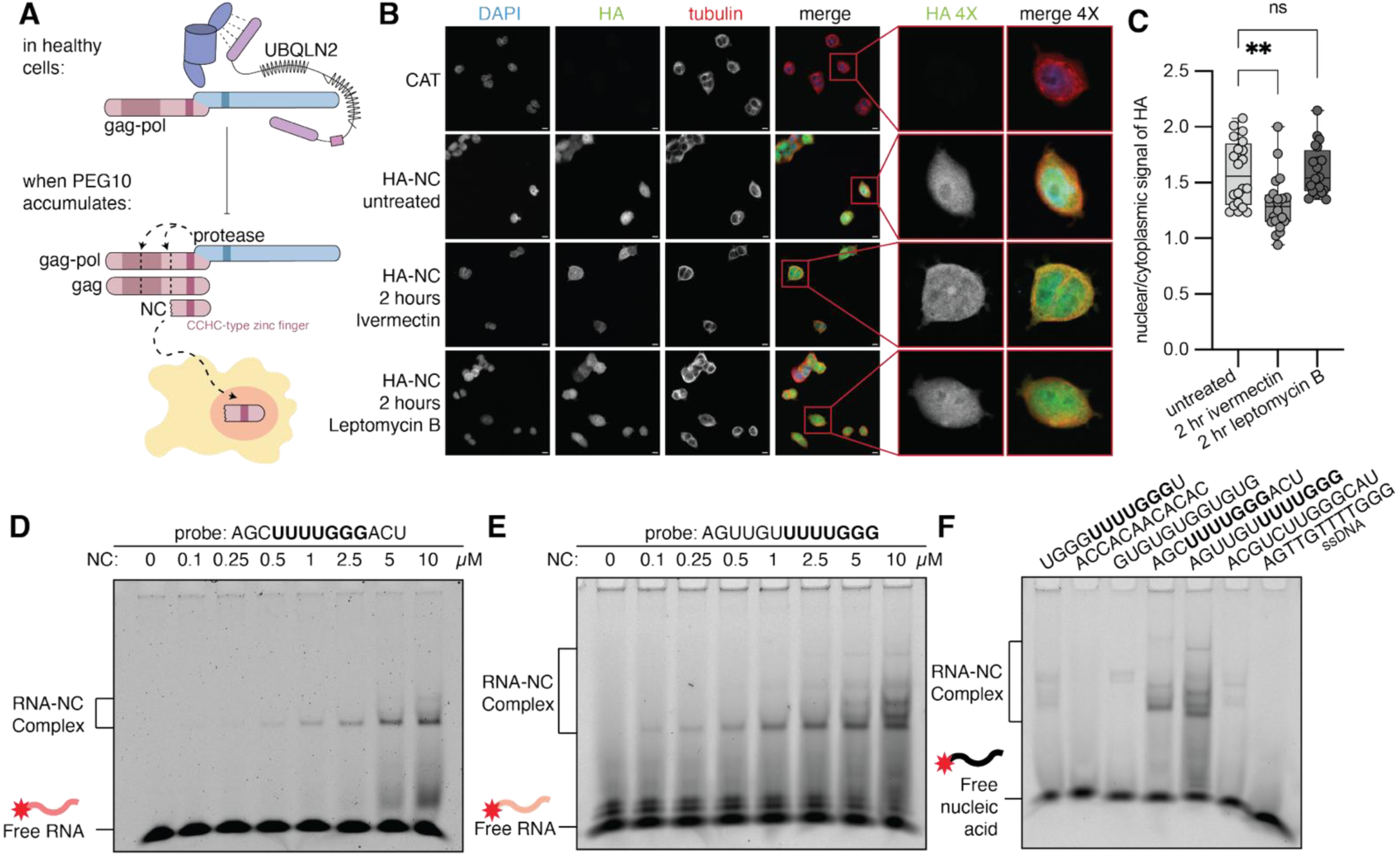
PEG10 NC binds ssRNA with a preference for a UUUUGGG motif. **(A)** Current, simplified model of PEG10 regulation, focusing on the cleavage and localization of the NC fragment during PEG10 self-processing. **(B)** IF staining of Flp-In-293 cells stably expressing a control of chloramphenicol acetyltransferase (CAT) or HA-NC comparing no drug treatment to 2 hours of ivermectin or leptomycin B treatment. Cells were stained with HA, DAPI, and tubulin. (n=2). On the right is 4x zoomed images of cells selected in red. **(C)** Quantification of the nuclear/cytoplasmic ratio of HA signal across drug treatments. (n=1) For statistical analysis, a one-way ANOVA was performed and compared to the nuclear/cytoplasmic ratio of untreated HA-NC cells. *p<0.05, **p<0.01, ***p<0.001, and ****p<0.0001. **(D-E)** fluorescent Electrophoretic Mobility Shift Assay (fEMSA) of PEG10 NC protein incubated with 0.5 nM Cy5.5-labeled RNA oligonucleotides. (n=3). NC-bound RNA shifts up in molecular weight. **(F)** fEMSA with 10 μM PEG10 NC incubated with 0.5 nM different Cy5.5-labeled oligonucleotides. (n=2).

## Results

### PEG10 NC traffics in and out of the nucleus

Previous work has demonstrated that the proteolytic NC fragment of PEG10 uniquely localizes to the nucleus, compared to other PEG10 constructs (Figure 1A)^23^. Many RNA-binding proteins, including those relevant to ALS such as TDP-43^40–42^, FUS^43,44^, and SFPQ^45^, traffic in and out of the nucleus to perform various cellular functions^46,47^. To evaluate the trafficking pattern of the NC fragment, we generated stable Flp-In-293 cell lines expressing HA-tagged NC protein (HA-NC) (Figure S1A). Cells were treated with ivermectin, to inhibit nuclear import^48^, or leptomycin B, to inhibit nuclear export^49^, and NC was visualized by immunofluorescence. At the steady state, more NC is found in the nucleus compared to the cytoplasm (Figure 1B-C). After 2 hours of treatment with ivermectin to inhibit active nuclear import, cells expressing HA-NC showed accumulation of HA signal in the cytoplasm of the cell (Figure 1B) with a marked decrease in the nuclear/cytoplasmic ratio (Figure 1C). This suggests that HA-NC trafficking into the nucleus is at least partially dependent on active import mechanisms. In contrast, 2 hours of inhibiting active nuclear export with leptomycin B had no effect on the localization of HA-NC compared to the steady state (Figure 1B-C), suggesting that either NC export from the nucleus is independent of active transport mechanisms, or that nuclear NC protein stays in the nucleus once delivered. Given the small size of NC (∼15 kDa with an HA tag), it is plausible that it may undergo passive transport if unbound to proteins or nucleic acids, but may otherwise rely on active nuclear transport.

### PEG10 NC demonstrates binding preference for U/G-rich RNAs

The CCHC-type zinc finger of NC is proposed to bind single-stranded nucleic acids, and in retrotransposons and retroviruses, the role of this zinc finger is to coordinate packaging of the RNA genome into capsids^50,51^. However, there has been disagreement about what nucleic acids PEG10 binds. It was first reported that PEG10 binds DNA^52^, but recent literature has suggested a variety of binding preferences, including bulk RNAs^53^, its own 3’ untranslated region (UTR)^54^, or a “UUUUGGG” (or, alternatively, a “UCUUGGG”) motif^55^. To explore PEG10’s nucleic acid binding preference, we performed fluorescent electrophoretic mobility shift assays (fEMSAs) with variations of the previously identified “UUUUGGG” motif to which PEG10 binds^55^. We observed purified NC protein binding to a “AGC**UUUUGGG**ACU” RNA probe (Figure 1D). We detected the highest level of binding for a “AGUUG**UUUUUGGG**” probe (Figure 1E), and some intermediate binding for a randomized U/G-rich RNA probe (Figure S1B) or a “UGGGG**UUUUGGG**U” RNA probe (Figure S1E). In contrast, NC never bound to A/C-rich RNA (Figure S1C), or a ssDNA probe of the best binding ssRNA (Figure S1D). However, NC was capable of binding the variation of the RNA motif, “ACG**U**C**UUGGG**CAU” (Figure S1F).

Then, each RNA probe was incubated with 10 μM NC and compared directly. We observed the highest level of binding to specific probes containing the “UUUUGGG” motif (Figure 1F), though there was considerable variability amongst motif-containing probes with unique flanking sequences, or the alternative motif including “**U**C**UUGGG**”. Together, our data confirms that NC has a preference for binding to U/G-rich RNA sequences, with less ability to bind to A/C rich sequences, and selectivity for RNA over DNA. Paired with evidence that NC traffics to the nucleus, this suggested a potentially broad role for PEG10 in regulating nuclear RNA.

### PEG10 does not mimic TDP-43 regulation of *STMN2*

While much remains unclear about the complex pathways involved in ALS disease progression, mutation or otherwise dysregulation of RNA binding proteins is commonly observed and is thought to directly contribute to neuronal dysfunction. Histological aggregates of TDP-43, a regulator of RNA splicing^56–59^, polyadenylation^58,60–63^, and mRNA transport^57,64–66^, are observed in 97% of ALS cases^5^. In addition, familial forms of ALS and ALS-FTD can be caused by mutation of the RNA binding proteins *TDP-43*^56,67^, *FUS*^68,69^, hnRNPs^70^, and others^71,72^. In many cases, a major contributor to RNA binding protein dysfunction is the mislocalization of these proteins. Given the U/G-rich RNA binding preference of NC, and its unique nuclear trafficking pattern upon gag-pol dependent protease cleavage, we investigated whether NC mimicked the regulatory activities of TDP-43.

To test whether NC could regulate splicing of a known TDP-43 splice target, *STMN2*^58,60,73^, we transfected SH-SY5Y cells, a neuroblastoma cell line, with HA-NC or *siTDP-43* and performed RT-PCR followed by agarose gel electrophoresis (Figure S2A-D). Knockdown of *TDP-43* with siRNA resulted in a relative accumulation of *STMN2* exon 2A (Figure S2D), confirming functional loss of TDP-43. In contrast, overexpression of NC did not significantly increase inclusion of exon 2A either by RT-PCR (Figure S2D), or qPCR (Figure S2E-F). To complement these findings, we also utilized a validated TDP-43 dependent splicing reporter construct where cryptic expression of an mScarlet reporter is revealed upon TDP-43 dysfunction^74^. While *TDP-43* depletion resulted in an expected observed increase of mScarlet expression (Figure S2G), we did not detect any reproducible changes to mScarlet expression upon overexpression of NC (Figure S2G). Therefore, we conclude that NC does not regulate *STMN2* akin to TDP-43, and decided to expand our focus to identifying neuronal transcripts that NC specifically regulates more broadly.

### PEG10 NC overexpression leads to widespread gene expression changes

Previous RNA-Sequencing (RNA-Seq) in HEK293 cells showed that overexpression of either gag-pol or NC protein increased transcripts in pathways of neuronal projection, axon extension, and axon guidance^23^. However, this experiment was limited by the use of a non-neuronal cell line^75^. To investigate gene expression changes in a model system more relevant to ALS, we performed RNA-Seq in SH-SY5Y cells following either depletion of *TDP-43*, depletion of *PEG10*, or overexpression of NC. We confirmed successful expression of HA-NC (Figure 2A) and a significant decrease in TDP-43 protein level (Figure 2B) and PEG10 protein level (Figure 2C) in our samples. *TARDBP* and *PEG10* were also depleted on the RNA level (Figure 2D-E). Global differential gene expression analysis identified thousands of genes which were differentially expressed upon either TDP-43 or PEG10 alteration (Figure 2F-H, Figure S3A-C, Table S1). Some changes to gene expression were shared between all conditions compared to *siNTC*, such as upregulation of *NNAT*, *SCAMP1*, *IGFBP5*, and *IQSEC1*, and downregulation of *TMEM160* (Figure 2F-H), implying that they may represent a generalized stress response to altered gene expression, or may provide a link between PEG10 and TDP-43 dysfunction. Other hits were shared between *PEG10* depletion and NC overexpression, such as depletion of *c4orf48* or enrichment of *BCYRN1* (Figure 2G-H), indicating that dysregulation of PEG10 in either direction may influence expression of particular genes. A similar phenomenon is observed for TDP-43, wherein overexpression and depletion of TDP-43 have both been shown to lead to RNA dysregulation^63,67,76,77^. Interestingly, NC overexpression led to more changes with log_2_FoldChange > 1 compared to all other conditions (Figure 2F-H). Many of these transcripts differ from previous RNA-Seq^23^ and enhanced crosslinking and immunoprecipitation (eCLIP) sequencing results^53^, suggesting that gene expression changes downstream of PEG10 modulation exhibit cell-type specificity.

**Figure 2.**
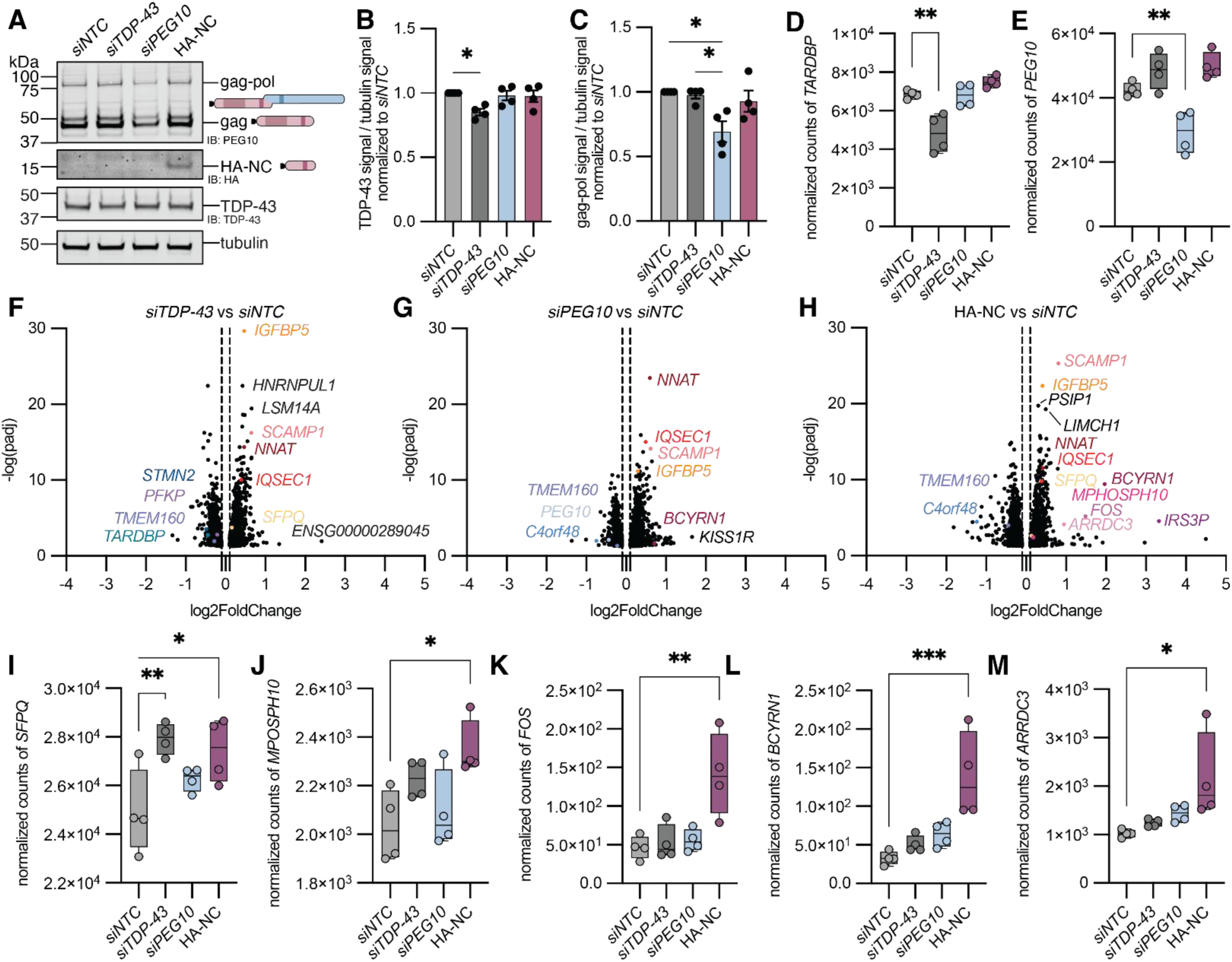
PEG10 NC regulates transcript abundance of genes involved in ALS. **(A)** Western blot of samples harvested in tandem with isolated RNA for RNA-Seq to confirm depletion of TDP-43 and *siPEG10*, and overexpression of HA-tagged NC (HA-NC). SH-SY5Y cells were transfected with *siNTC*, *siTDP-43*, *siPEG10*, or HA-NC for 48 hours prior to harvesting, then probed by western blot for endogenous PEG10, HA, TDP-43, and tubulin. (n=4). **(B-C)** Quantification of western blot results in A for **(B)** TDP-43 and **(C)** PEG10 gag-pol protein. (n=4). **(D-E)** Normalized counts of **(D)** *TARDBP* and **(E)** *PEG10* RNA from the RNA-Seq experiment comparing *siNTC, siTDP-43, siPEG10,* and HA-NC. For (B-E), a one-way ANOVA was performed and compared to the mean of siNTC. **(F-H)** Volcano plots of RNA-Sequencing results of normalized counts comparing **(F)** *siTDP-43***, (G)** *siPEG10* or **(H)** HA-NC overexpression to *siNTC* as a negative control. Some top hits are named and color coded for specific genes that are shared between conditions. (n=4). **(I-M)** Normalized counts of **(I)** *MPHOSPH10***, (J)** *FOS***, (K)** *BCYRN1***, (L)** *ARRDC3*, and **(M)** *SFPQ* transcripts across each condition (n=4). For (F-M), statistics were determined by one-way ANOVA. *p<0.05, **p<0.01, ***p<0.001, and ****p<0.0001.

### RNAs implicated in neurodegenerative diseases are altered upon NC overexpression

NC overexpression caused changes to genes that are implicated in neurodegenerative disease. Expression of *splicing factor proline- and glutamine-rich (SFPQ),* a splicing factor which is elevated in the cytoplasm in ALS^45,78^ and can cause fALS when mutated^79^, was increased upon both *TDP-43* depletion and NC overexpression (Figure 2I). Expression of *FOS,* a transcription factor increased in *FUS*-ALS that contributes to aberrant axon branching in motor neurons^80^, was elevated specifically upon NC overexpression (Figure 2J). NC overexpression also induced gene expression changes that have not been explicitly linked to ALS but have been investigated in other neurodegenerative conditions. *Brain cytoplasmic RNA 1* (*BCYRN1,* also known as *BC200*) is a long non-coding RNA (lncRNA) elevated in Alzheimer’s Disease (AD) ^81^ and was significantly accumulated in NC overexpressing cells (Figure 2K). NC overexpression also resulted in an accumulation of *Arrestin Domain Containing 3* (*ARRDC3*) (Figure 2L), which is also elevated in AD organoids^82^.

### NC overexpression mimics TDP-43 loss in pathways of gene regulation

In addition to looking at changes to normalized counts of RNAs differentially expressed in each condition, we also looked at trends in gene expression changes through analysis of Gene Ontology (GO)-term enrichments^83^ across each condition (Figure 3A-C, Figure S3D-I, Table S1). Upon *TDP-43* depletion, there was overrepresentation of mRNA processing and mRNA splicing pathways (Figure 3A), which is consistent with TDP-43’s central role in RNA biology^41,64,84^. *PEG10* depletion led to overrepresentation of pathways involved in signaling, axon development, and axonogenesis (Figure 3B). Changes to these pathways were unexpected, as similar upregulation of axon extension and maintenance pathways were observed upon NC overexpression in previous RNA-Seq experiments^23^. However, the most striking finding was that NC overexpression, like *TDP-43* knockdown, resulted in the upregulation of mRNA processing and RNA splicing pathways (Figure 3C). This data demonstrates a role of NC in regulating expression of RNA splicing factors, which has not been previously described.

**Figure 3.**
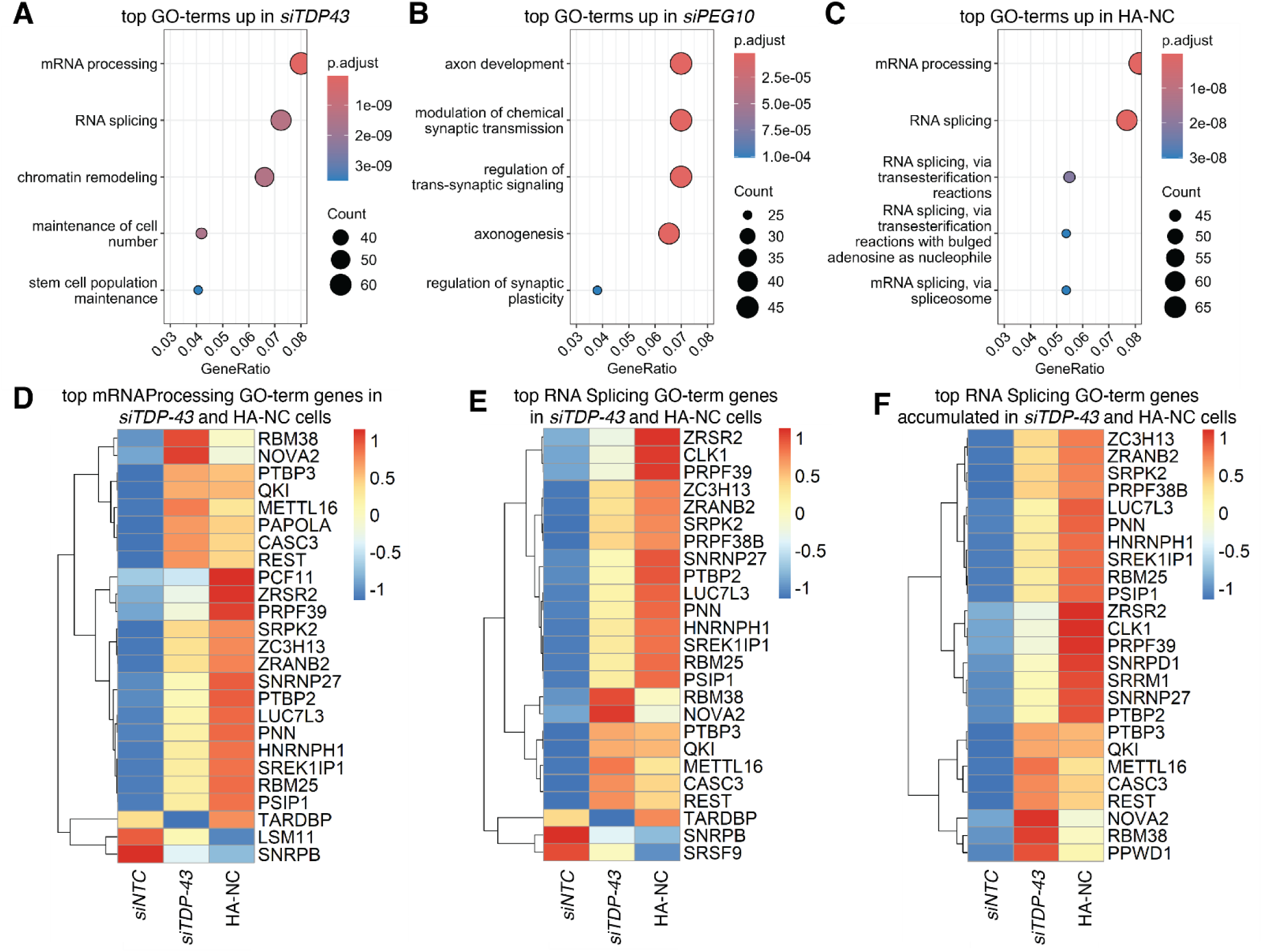
PEG10 NC overexpression mimics and exacerbates TDP-43 depletion. **(A-C)** Top upregulated pathways of gene expression changes in cells transfected with **(A)** *siTDP-43,* **(B)** *siPEG10*, or **(C)** HA-NC by Gene Ontology (GO)-term enrichment analysis. The top five GO-terms using a log_2_FoldChange cutoff of 0.1 were ranked by adjusted p-value. Adjusted p-value is shown by color, and the size of the datapoint reflects the number of genes enriched in the pathway. **(D)** Heatmap of genes from the mRNA processing GO-term showing with color scale indicating the z-score for the top 25 differentially expressed genes ranked top to bottom by log_2_FoldChange**. (E)** Heatmap of genes from the RNA splicing GO-term with color scale indicating the row z-score for the top 25 differentially expressed genes in either direction ranked top to bottom by log_2_FoldChange. **(F)** Heatmap of genes from the RNA splicing GO-term showing row z-score for the top 25 accumulated genes ranked top to bottom by log_2_FoldChange. This demonstrates that most changes to splicing GO-terms are upregulation, as E and F show very similar top terms. In D-F, only changes with l log_2_FoldChange greater than 0.1 were evaluated. (n=4).

Closer examination of RNA splicing-related GO-terms revealed both shared and unique changes upon *TDP-43* depletion or NC overexpression (Figure 3D-F), with some genes like *PTPB3* and *QKI,* in particular, being shared between TDP-43 and NC dysregulation (Figure 3D-F). The enrichment of mRNA splicing factors upon NC overexpression suggested that like TDP-43, NC dysregulation could be deranging mRNA splicing, leading to upregulation of splicing factors in response. Therefore, our next step was to more directly investigate how mRNA splicing changed upon NC overexpression.

### PEG10 NC regulates splicing of *NRG3* in neuronal cells

MAJIQ analysis^85^ was then performed to evaluate specific changes to mRNA splicing. After generating a list of genes with significantly altered splicing for each condition compared to *siNTC* (Table S2), we generated a Venn Diagram of splice changes for each RNA with observed changes to local splice variations (Figure 4A). As expected, *STMN2* splicing was significantly altered by knockdown of *TDP-43* (Figure 4A). Counts of *STMN2* mRNA were significantly depleted (Figure 4B), and *STMN2* cryptic exon 2A is present only in *siTDP-43* treated cells (Figure 4C). Consistent with our findings from *STMN2* RT-PCR and reporter experiments (Figure S2), there was no change to *STMN2* counts or inclusion of *STMN2* exon 2A upon PEG10 dysregulation, further highlighting that the specific effects on mRNA splicing caused by TDP-43 disruption and PEG10 accumulation are distinct.

**Figure 4.**
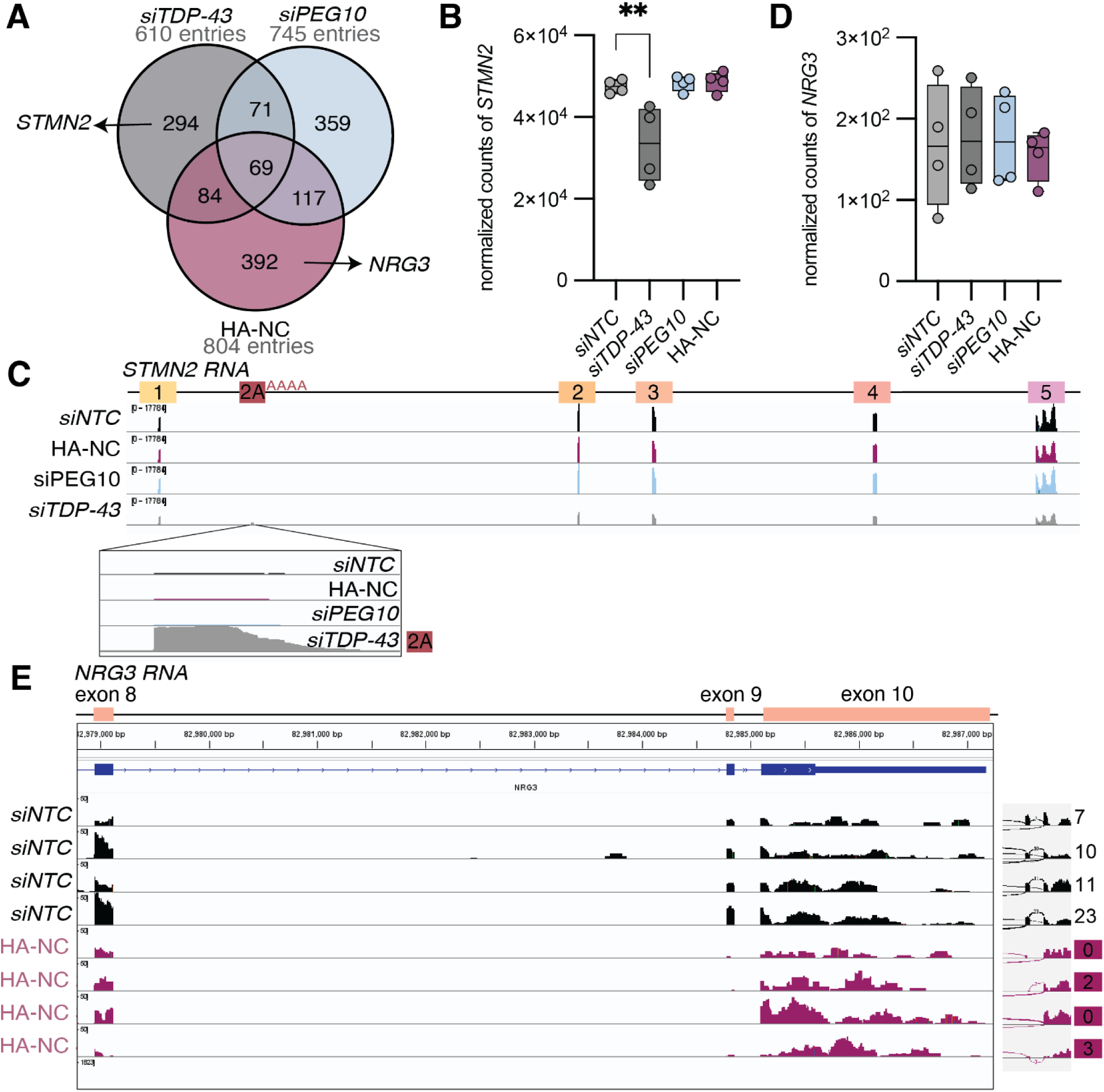
PEG10 NC causes distinct splicing changes to TDP-43. **(A)** Venn diagram highlighting splice changes among *siTDP-43*, *siPEG10*, and HA-NC transfected cells using MAJIQ with Voila viewer. Splice changes were determined using a deltapsi cutoff of 0.1 and a confidence interval cutoff of 0.9. **(B)** Normalized counts of full-length *STMN2* transcript across conditions, showing decreased abundance of full-length *STMN2* in siTDP-43 expressing cells. (n=4). **(C)** Representative IGV snapshot of *STMN2* across each condition of RNA-Seq experiment, highlighting the cryptic exon splice event of *STMN2* exon 2A for *siTDP-43*. Schematic of *STMN2* mRNA with numbered exons shown at the top. At bottom is a zoomed in region with exon 2A. **(D)** Normalized counts of full-length *NRG3* transcript across conditions. (n=4). For (B,D), statistics were determined by DESeq2. **(E, left)** IGV snapshot of the transcript counts of *NRG3.* Snapshot view includes IGV location data for exons 8-10 of *NRG3*, showing that exon 9 is skipped in HA-NC cells compared to WT. Schematic of *NRG3* mRNA with numbered exons shown at the top. **(E, right)** IGV snapshot of the Sashimi plot of a portion of *NRG3* for all four biological replicates of *siNTC* compared to HA-NC transfected cells. This view shows the exon skipping event identified by MAJIQ.

One aberrant alternative splicing change observed only upon NC overexpression was an exon skipping event in *neuregulin 3 (NRG3)* (Table S2)*. NRG3* encodes an epidermal growth factor (EGF)-like signaling molecule that is expressed in the brain^86^. The NRG3 precursor protein is cleaved by BACE1 to generate an EGF-like ligand that binds to and stimulates the Erb-B2 receptor tyrosine kinase 4 (ErbB4), leading to tyrosine phosphorylation of the receptor and signal transduction^86–89^. The neuregulin-ErbB4 pathway influences many neuronal processes, including synaptic development, neuronal proliferation, and neurotransmission^90^. Additionally, dysregulation of the neuregulin-ErbB4 pathway via mutation of *ERBB4* causes a rare form of fALS^91^. The *NRG3* pre-mRNA includes 10 exons, spans 1.1 Mb and undergoes extensive alternative splicing to generate multiple isoforms of mature *NRG3* transcript^86,88,92^. MAJIQ analysis revealed little or no inclusion of exon 9 in mature mRNA of NC-expressing cells, which would result in an in-frame loss of 72 nucleotides. There were no changes to *NRG3* RNA abundance (Figure 4D), suggesting that this splice change is unlikely to result in significant nonsense-mediated decay. We confirmed these splicing changes by transcript counts per exon in IGV, where we see decreased or no counts at exon 9 in SH-SY5Y cells expressing NC protein (Figure 4E).

### *NRG3* splicing changes lead to decreased protein abundance in SH-SY5Y cells

The exon skipping event observed for *NRG3* leads to a 24 amino acid in-frame deletion in the cytoplasmic domain of NRG3 (Figure 5A). While this exon skipping event has been annotated^92^, it has not been reliably observed or characterized, and the functional consequence of this deletion remains unknown. While the cytoplasmic domain is not directly involved in receptor binding to ErbB4, it is proposed to regulate trafficking and proteolytic processing of NRG3^93^. Therefore, an exon skipping event in this domain could have detrimental effects on protein folding, trafficking, proteolytic processing, and stability of the NRG3 protein.

**Figure 5.**
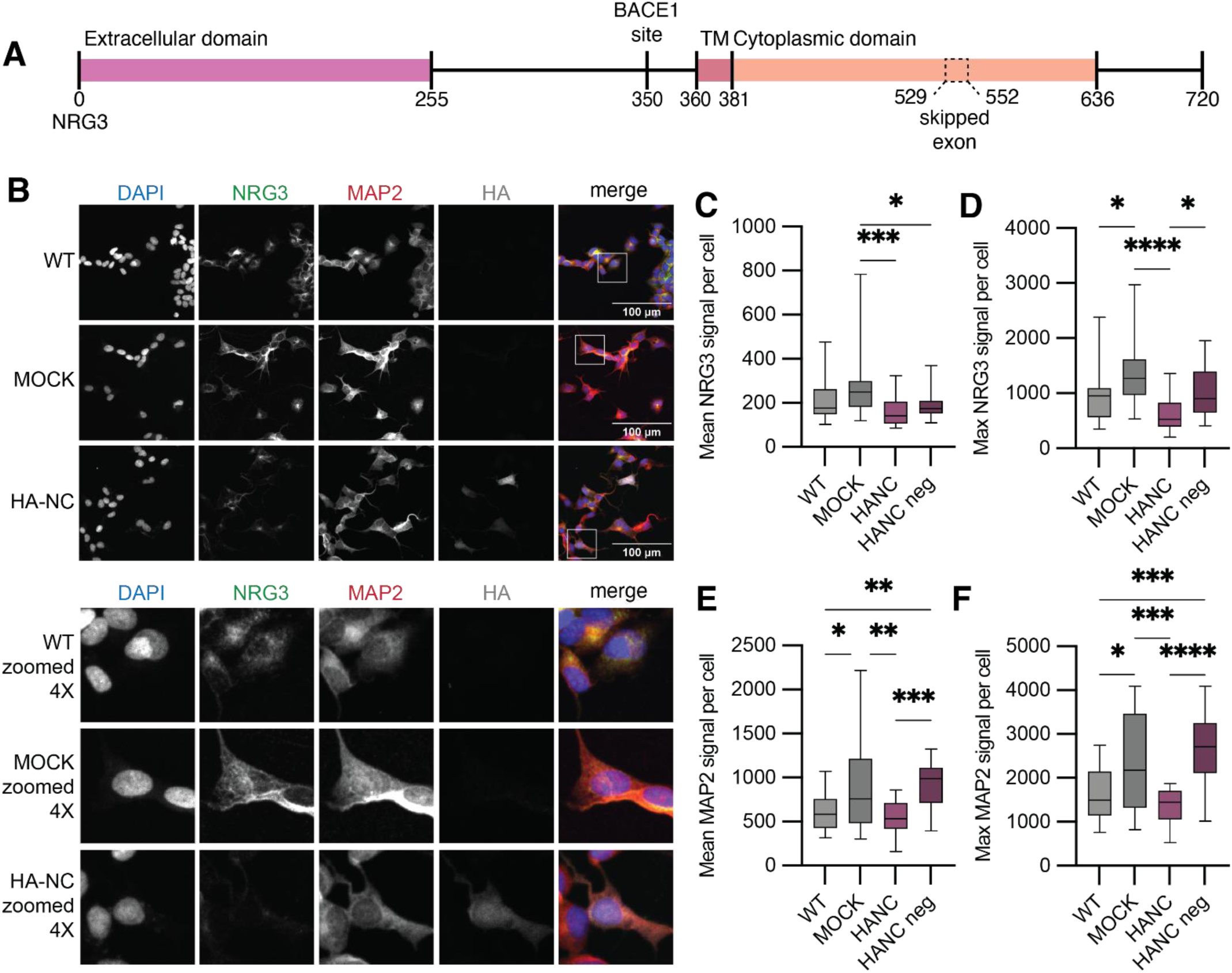
Expression of PEG10 NC leads to lower NRG3 staining in SH-SY5Y cells. **(A)** Simplified schematic of NRG3 protein, illustrating the extracellular domain, BACE1 cleavage site, transmembrane domain (TM), and the cytoplasmic domain. Amino acid numbers are shown at bottom. The predicted exon skipping event is shown in dotted lines to demonstrate relative location of the skipped exon in the cytoplasmic domain. **(B, top)** Representative images from IF staining of WT SH-SY5Y cells compared to MOCK or HA-NC transfected cells. Cells were stained with NRG3, MAP2, and HA and coverslips were mounted with Prolong Gold with DAPI. Z-stack images were collected and scale bar = 10 μm. (n=3). **(B, bottom)** 4X zoomed images of cells in F, highlighting example cells for each condition. **(C)** Mean NRG3 intensity across Z-stacks for single cells in each condition for one representative experiment. HA-positive and HA-negative cells from the same coverslip were compared in purple. **(D)** Maximum NRG3 signal across Z-stacks for single cells in each condition for one representative experiment. **(E)** Mean MAP2 intensity across Z-stacks for single cells in each condition. **(F)** Maximum MAP2 signal intensity across Z-stacks for single cells in each condition. For (C-F), n=20-22 cells per condition, and a one-way ANOVA was performed and compared to each mean. *p<0.05, **p<0.01, ***p<0.001, and ****p<0.0001.

To evaluate NRG3 protein abundance and localization in cells expressing PEG10 NC, we performed IF staining of NC-transfected cells using an antibody that recognizes an epitope from AA 560-720 of the NRG3 protein (Figure 5B). Both the mean and maximum NRG3 signal per cell was decreased in HA positive cells compared to mock transfected cells (Figure 5C-D), indicating that expression of NC protein leads to changes in NRG3 expression. When we quantified the NRG3 signal compared to MAP2 as a control, we noticed that MAP2 was also significantly decreased in NC expressing cells (Figure 5E-F). However, to account for variability between coverslips, we compared NRG3 and MAP2 signal between HA positive and HA negative cells on the same coverslip and observed less NRG3 and MAP2 signal for HA positive cells (Figure 5C-F). Whole cell lysate of transfected SH-SY5Y cells showed no observable changes in molecular weight, or the overall abundance of NRG3 (Figure S4A-B); however, it is possible that untransfected cells masked the effects of NC transfection in this bulk analysis (Figure S4C).

### NC expression in human iNeurons leads to decreased dendritic NRG3

To identify consequences of *NRG3* exon 9 skipping in human neurons, we utilized WT and NC expressing NGN2-inducible human embryonic stem cells (hESCs) to generate induced neurons (iNeurons)^94^. We performed IF staining of differentiated WT and HA-NC iNeurons to evaluate changes to morphology and NRG3 expression (Figure 6A-B). Sholl analysis^95,96^ revealed that NC-expressing iNeurons were more complex than WT iNeurons (Figure 6C). In addition to increased complexity, primary and secondary dendrites were longer overall for NC-expressing iNeurons (Figure 6D-E).

**Figure 6.**
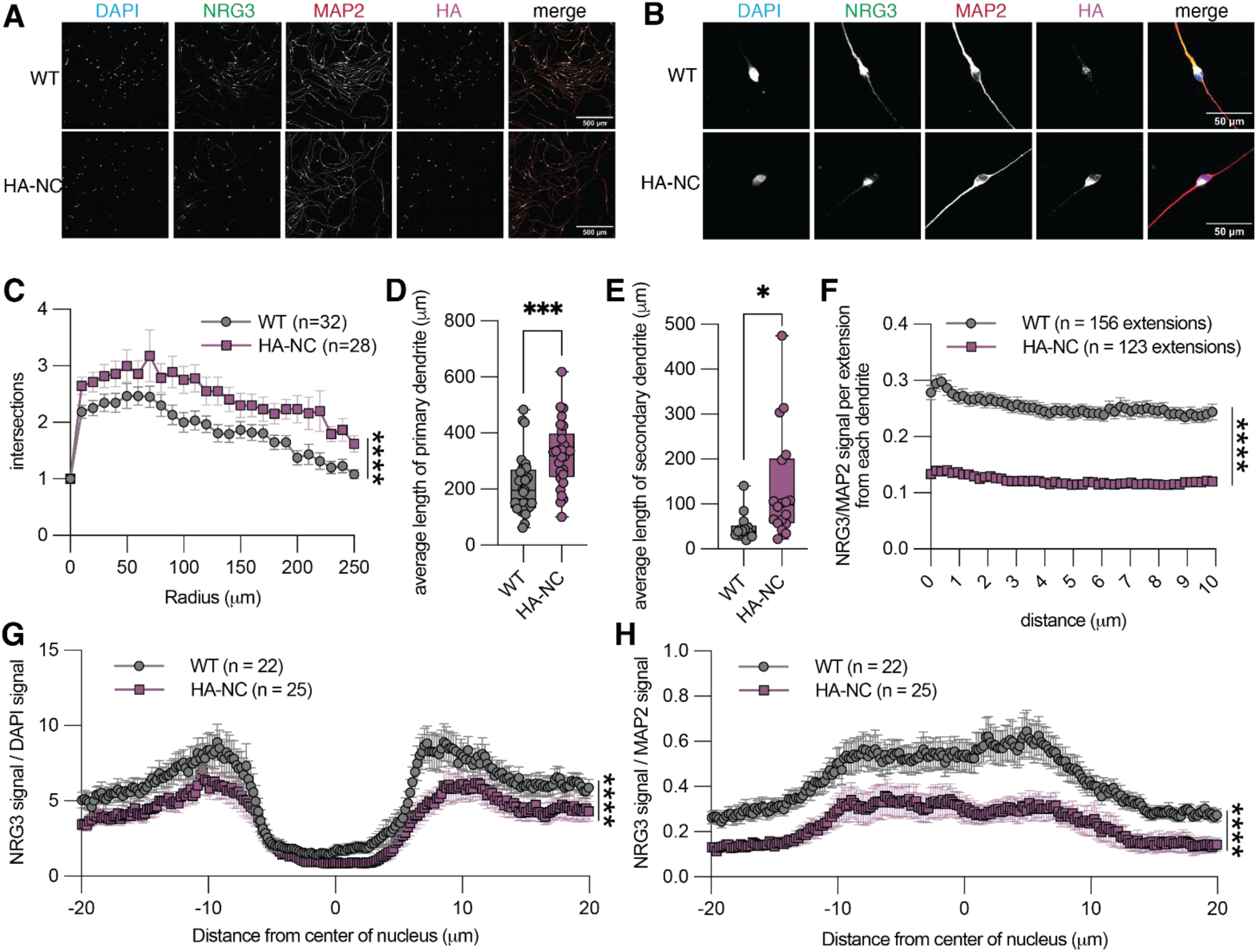
NRG3 is depleted along dendrites in PEG10 NC expressing iNeurons. **(A-B)** Representative IF images of WT and HA-NC iNeurons differentiated from hESCs for 7 days and then stained for NRG3, MAP2, and HA (n=3) at **(A)** 20X, with scale bar of 100 μm, or **(B)** 100X, with scale bar of 50 μm. Contrast was enhanced for (A-B) for ease of visualization. **(C)** Sholl analysis of WT and HA-NC iNeurons (n=2 independent experiments, 20 – 32 iNeurons per condition per experiment). Statistics were determined via two-way ANOVA between WT and HA-NC iNeurons. **(D – E)** average length of **(D)** primary and **(E)** secondary dendrites per iNeuron in C. For statistical analysis in (D-E), a Student’s t-test was performed. **(F)** mean NRG3/MAP2 signal from dendrites starting at the branch point from the soma (n=3 independent experiments, n=59 – 156 iNeurons per condition per experiment). Statistics were determined via two-way ANOVA between WT and HA-NC iNeurons. **(G-H)** NRG3 signal over **(G)** DAPI or **(H)** MAP2 starting at the nucleus (0) and extending 20 micrometers in either direction (−20 and 20) across primary dendrites extending from the soma (n=1 quantified experiment, representative of n=3 experiments from F). Statistics were determined via two-way ANOVA between WT and HA-NC iNeurons *p<0.05, **p<0.01, ***p<0.001, and ****p<0.0001.

To evaluate NRG3 abundance across iNeurons, we tracked the NRG3/MAP2 signal along primary dendritic branches extending from the soma. We observed a consistent and significant decrease in NRG3/MAP2 signal across dendrites when NC is expressed (Figure 6F). Unlike SH-SY5Y cells, iNeuron MAP2 was not significantly influenced by NC expression (Figure S5A-B). To determine whether the loss of NRG3 was specific to dendrites, we also tracked the NRG3 signal normalized to either DAPI or MAP2 aling a line profile spanning the iNeuron and observed a decrease in NRG3 across the whole of the iNeuron (Figure 6G-H). Based on our data, we conclude that NRG3 is reduced in iNeurons expressing NC, likely due to the exon skipping event of *NRG3* exon 9.

### *NRG3* undergoes exon skipping in cases of ALS

Results presented thus far provide compelling evidence that NC-mediated NRG3 mis-splicing leads to decreased protein levels in human iNeurons, but the direct link to disease remains elusive. To address this, we explored previously analyzed ALS patient gene expression data^23^. Some of the genes most altered by NC overexpression, including *c4orf48*, *Fragile Messenger Ribonucleoprotein 1 (FMR1),* and *PYGM*, are also significantly altered in postmortem lumbar spinal cord samples from ALS patients (Figure 7A-F). We did not observe a decrease of *NRG3* in ALS (Figure 7G); however, *NRG3* normalized counts were not altered in SH-SY5Y cells upon NC overexpression (Figure 4D).

**Figure 7.**
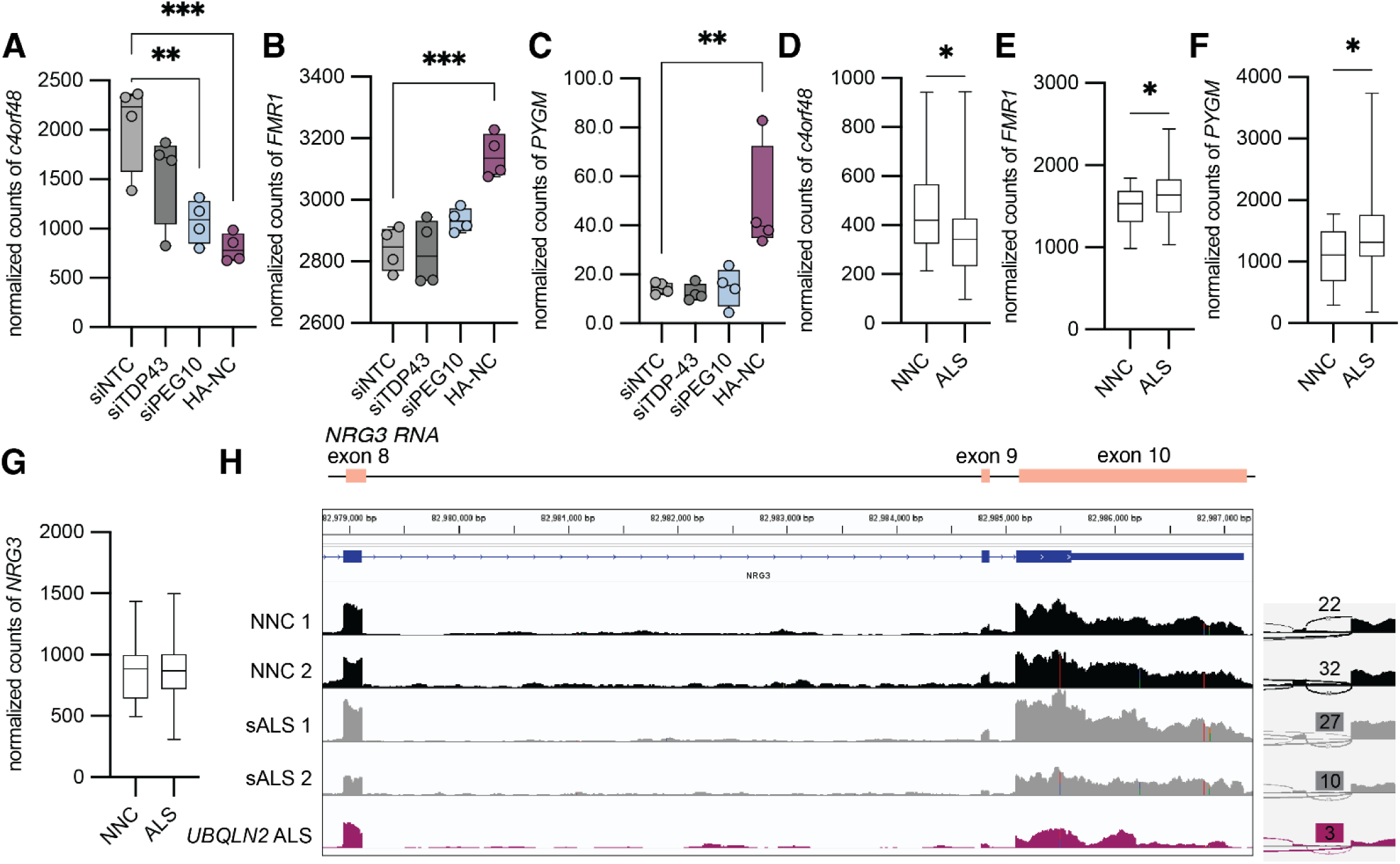
PEG10 NC influences gene expression of transcripts significantly altered in ALS. **(A-C)** Normalized counts of **(A)** *c4orf48* **(B)** *FMR1,* and **(C)** *PYGM* from RNA-Seq of transfected SH-SY5Y cells. For statistical analysis in (A-C), a one-way ANOVA was performed and compared to the mean of *siNTC*. **(D-G)** Normalized counts of **(D)** *c4orf48* **(E)** *FMR1,* **(F)** *PYGM,* and **(G)** *NRG3* using RNA-seq data from the Target Amyotrophic Lateral Sclerosis dataset of post-mortem lumbar spinal cords. n=16 NNC and n=126 ALS samples. For statistical analysis in D-G, a Student’s t-test was performed. **(H, left)** IGV snapshot of the transcript counts from RNA-seq data from the Target Amyotrophic Lateral Sclerosis dataset of post-mortem lumbar spinal cords. This view includes exons 8 – 10 of *NRG3*, showing that exon 9 is skipped in *UBQLN2*-mediated ALS and case 2 of sALS. **(H, right)** IGV snapshot of the Sashimi plot of a portion of *NRG3* for the samples on the left, showing counts of exon 9 inclusion. NNC = non-neurological control. *p<0.05, **p<0.01, ***p<0.001, and ****p<0.0001.

While PEG10 gag-pol levels were elevated in a small cohort of lumbosacral spinal cord from sALS patients^23^, it is unclear whether elevated PEG10 is expected to be a universal finding in sporadic disease, which dominates the dataset we queried. Therefore, we might expect that some NC-induced gene expression changes observed in SH-SY5Y cells independently mimic gene expression changes observed in sALS, while other gene expression changes shared between NC overexpression and ALS tissue samples could reflect PEG10-dependent pathology in samples where PEG10 is elevated. As one example of an ALS case with the clearest connection to elevated PEG10, we age and sex matched the single available *UBQLN2*-mediated ALS case with 2 non-neurological controls (NNC) and two sporadic ALS cases, and investigated *NRG3* exon 9 splicing. We observed a reduction in *NRG3* exon 9 in human *UBQLN2*-mediated ALS, and a moderate reduction in one of the sALS cases (Figure 7H). When we compared other transcript changes, we also observed that both the *UBQLN2*-mediated ALS case and the sALS case with decreased *NRG3* exon 9 also had increased levels of *BCYRN1*, similar to what was observed with SH-SY5Y RNA-Seq results (Figure S5C-D, Figure 2L). Therefore, we would predict that the sALS case with increased *BCYRN1* and decreased *NRG3* exon 9 has increased protein abundance of PEG10. These findings demonstrate a novel role for the retrotransposon-derived protein PEG10 in regulating mRNA splicing in neurons and suggest that these splicing changes influence neuronal health in ways that mimic the pathology of ALS. Paired with recent evidence that PEG10 levels are elevated in ALS-FTD, these results suggest that dysregulation of PEG10 may contribute to neuronal dysfunction in a mechanism akin to, but distinct from, TDP-43.

## Discussion

PEG10 is a domesticated retroelement that resembles the gag-pol polyprotein of retroviruses and retrotransposons^32,36^ that has been linked to ALS-FTD^22,23^ and Angelman’s syndrome^24^ through unbiased screening techniques, and may contribute to pathology by influencing gene expression. As part of its unique self-regulation, the protease of the pol region directs cleavage and liberation of a nucleic-acid binding proteolytic fragment called ‘NC’ that moves to the nucleus and directs gene expression changes in the cell^23^. This work focused on evaluating the specific effects of NC accumulation in multiple cellular models, including human neurons.

We first showed that NC traffics to the nucleus and has a preference for interacting with U/G-rich RNA, settling confusion as to the binding preference of this protein for nucleic acid. This highlighted similarities between NC and TDP-43: both are nuclear nucleic acid binding proteins that bind U/G-rich RNAs, and both are altered in ALS^23,41,64,97,98^. Notably, while TDP-43 is mislocalized from the nucleus to the cytoplasm in ALS^41^, the accumulation of PEG10 protein, through mutation or aggregation of its proteolytic regulator UBQLN2, results in NC proteolytic cleavage and an increase in nuclear localization in disease conditions.

Utilizing RNA-Seq, we identified specific transcripts that are altered upon overexpression of NC or depletion of *PEG10* or *TDP-43*. We observed both shared and distinct transcript changes between knockdown and overexpression, highlighting that modulation of PEG10 in either direction contributes to changes in gene expression. This was surprising, but little is known about the effects of PEG10 depletion outside of the reproductive system. In mice, *PEG10* deletion, protease mutation, or targeted depletion of gag-pol protein results in embryonic or perinatal lethality due to placentation defects^99–101^. Molecular studies suggest that in placental precursor cells, PEG10 modulates expression of genes involved in trophoblast differentiation and survival^53^, but future work is required to determine if PEG10 plays a parallel role in the nervous system. At the level of pathways of gene expression, in contrast, NC overexpression strongly mimicked knockdown of *TDP-43* with enrichment for pathways of mRNA splicing and RNA regulation.

Despite shared GO-term enrichment pathways, NC overexpression and TDP-43 depletion led to distinct splicing changes. Unlike TDP-43, we did not observe any particular enrichment for cryptic exon inclusion with NC overexpression. This may reflect that overexpression of NC leads to increased occupancy in the nucleus, and presumably more saturation of NC on U/G-rich RNA sequences, as opposed to a depletion of TDP-43 from nuclear pre-mRNA, leading to cryptic exon splicing in its absence. Further, we did not observe changes to classical TDP-43 target transcripts, including *STMN2*, either by RT-PCR, reporter, or RNA-Seq, indicating that NC overexpression does not likely interfere with TDP-43 activity. This is further supported by the apparent preference of NC for U/G rich sequences, as opposed to the UG-rich repeats preferred by TDP-43^74,98^, making it likely that these two proteins have unique mRNA binding profiles.

We determined that NC regulates splicing of *NRG3*, an neural-enriched signaling protein^86–89^. The overexpression of NC resulted in a near-complete exclusion of exon 9, which results in an in-frame deletion of a portion of the cytoplasmic domain thought to regulate intracellular receptor trafficking^93^. Consistent with this hypothesis, we found that in both SH-SY5Y cells and human neurons expressing NC, there was a decrease in NRG3 protein, suggesting that this splicing defect leads to changes in protein abundance or localization of NRG3. NRG3 signals through ERBB4, which has independently been linked to familial ALS^91^, providing a potential link between PEG10, aberrant splicing, and ALS. Future work is required, however, to determine if PEG10-dependent NRG3 splicing changes are a direct consequence of NC binding to *NRG3* transcript or are an indirect consequence of the dysregulation of other splicing factors as observed by pathway enrichment.

Morphological studies of iNeurons concluded that NC expression leads to increased dendritic complexity and longer dendrites. In contrast, *UBQLN2* mutations, which lead to elevated PEG10 levels, result in decreased dendritic complexity^102,103^. One possibility is that acute PEG10 expression leads to an initial, yet unsustainable, burst of dendritic branching, perhaps due to changes in gene expression such as *FOS*^80^. Another possibility is that the genetic mutation of *UBQLN2* leads to chronic PEG10 levels in precursor cells that influence neuronal differentiation in ways distinct from our experimental approach with NC expression driven by the Synapsin promoter. Finally, PEG10 is one of many changes caused by *UBQLN2* mutations^22,23,31,104,105^, and while PEG10 accumulation may induce dendritic branching, there may be other changes that result in overall spinopathy in cases of *UBQLN2*-mediated disease. Future work is needed to examine these possibilities.

In postmortem RNA-Seq data from human patients with ALS, we observed decreased *NRG3* splicing of exon 9 in *UBQLN2*-mediated ALS compared to non-neurological controls. Based on our human data, we expect that this splicing defect leads to decreased NRG3 and, ultimately, signaling capacity in ALS neurons when PEG10 is accumulated. A challenge of interpreting this data, however, is that it is challenging to predict which samples might exhibit PEG10-dependent transcriptional changes when the protein levels of PEG10 are unknown for most samples. The one available *UBQLN2*-mediated ALS case has high levels of PEG10 gag-pol protein^23^, and a small cohort of sporadic cases has shown elevated PEG10 compared to control^23^, but it is unknown if this is universal in ALS. We predict that cases with increased *BCYRN1* and the *NRG3* exon skipping event have higher PEG10 protein abundance and could act as transcriptional biomarkers for PEG10 accumulation in ALS.

Overall, we have defined a novel role for PEG10, through the proteolytic generation of NC protein, in regulating mRNA splicing and gene expression in human neurons in a manner reminiscent of, but distinct from, TDP-43. We show data that suggests that these changes also occur in ALS patients, thereby providing a compelling connection between PEG10 accumulation and ALS disease pathology. Far from being an evolutionary fossil, PEG10 is independently capable of dysregulating RNA splicing in human neurons, making it a new addition to the library of RNA binding proteins linked to ALS-FTD and a likely contributor to disease.

## Resource availability

### Lead contact

Further information and requests for resources and reagents should be directed to Dr. Alexandra Whiteley.

### Materials availability

Plasmids generated for this work are available upon request from the lead contact.

### Data and code availability

Raw data will be shared by the lead contact upon request. Any additional information and code required to reanalyze the data reported in this paper is available from the lead contact upon request.

## Acknowledgments

We sincerely thank Phuoc Huyhn, Maria Rogan, Kian Grimison, Katie Waldon, and past lab members of the Whiteley laboratory, including G. Aaron Holling and Julia Roberts, for helpful discussion and comments. We would also like to acknowledge the Target ALS Human Postmortem Tissue Core for post-mortem RNA-Seq reads. We want to give a special thank you to Will Campodonico for originally generating the Bam files and FeatureCounts from the TargetALS dataset, and for providing helpful critical feedback on the manuscript. This work was supported by the NIH NIGMS Signaling and Cellular Regulation Graduate Training Program (T32GM142607) to A.M. Matthews and NIH NINDS R01 NS131660 to A.M. Whiteley. We would also like to acknowledge the Shared Instruments Pool (SIP) through the Department of Biochemistry at CU Boulder (RRID: SCR_018986), the Biochemistry Cell Culture Core Facility (CCF) (RRID: SCR_018988), and the BioFrontier Institute’s Advanced Light Microscopy Core (ALMC) (RRID: SCR_018302) for their assistance with shared equipment. Laser scanning confocal microscopy was performed on a Nikon A1R microscope supported by NIST-CU Cooperative Agreement award number 70NANB15H226 or the Nikon AXR Laser Scanning Confocal, which is supported by NIH Grant 1S10OD034320. We would like to give a special thank you to Dr. Joseph Dragavon, Dr. Jian Wei Tay, and Dr. Erin Richards through the ALMC for their amazing technical assistance. We would also like to thank Theresa Nahreini and Emily Proksch of the CCF and Dr. Annette Erbse for their invaluable help in utilizing shared equipment and reagents. We would also like to thank Drs. Roy Parker, Sabrina Spencer, Kelsie Eichel, and Edward Chuong for their critical evaluation of this work.

## Author contributions

conceptualization: A.M. Matthews and A.M. Whiteley; experimentation: A.M. Matthews; data analysis: A.M. Matthews and A.M. Whiteley; writing – original draft: A.M. Matthews; writing – review and editing: A.M. Matthews and A.M. Whiteley; funding acquisition: A.M. Whiteley.

## Materials and Methods

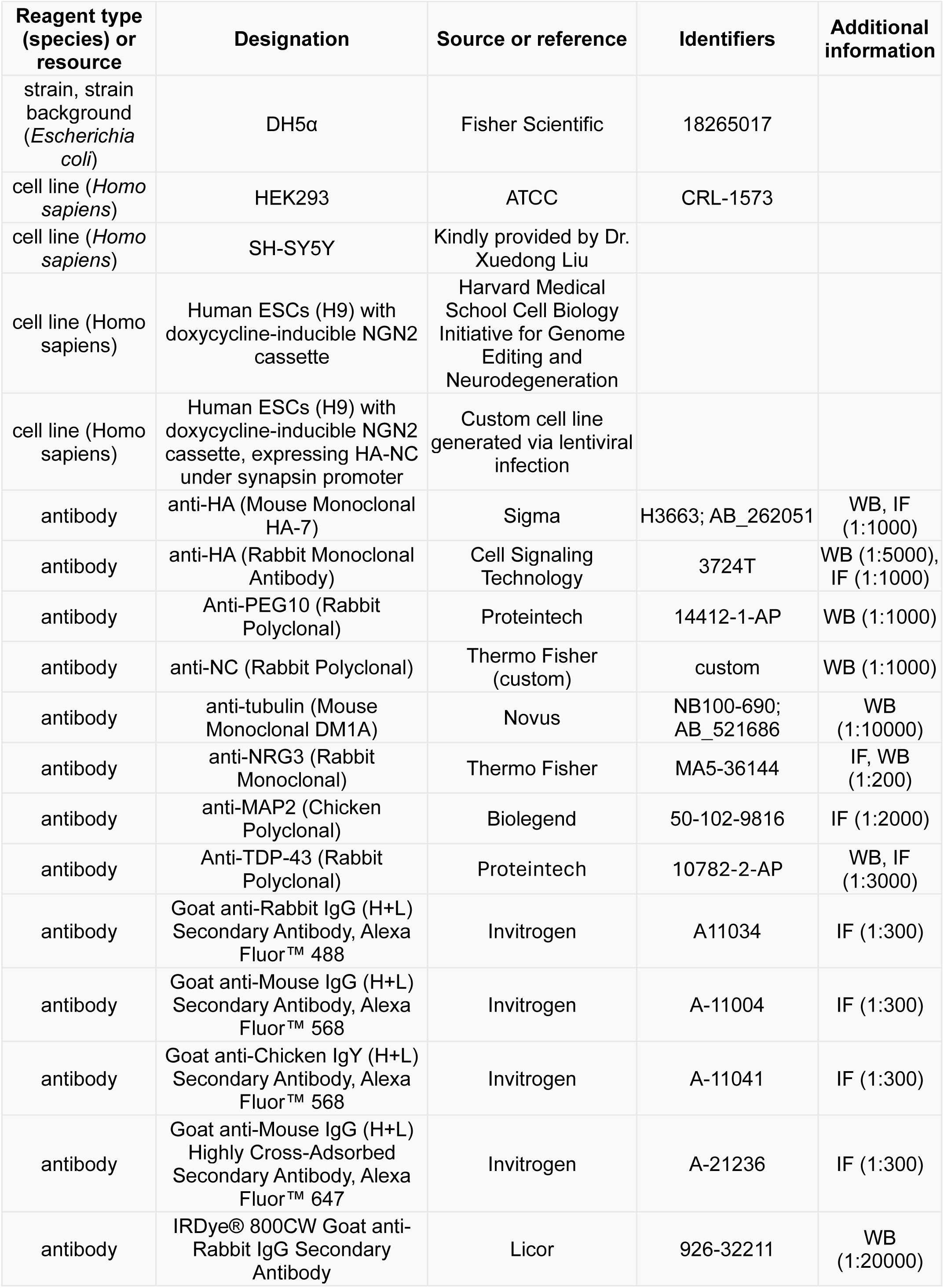
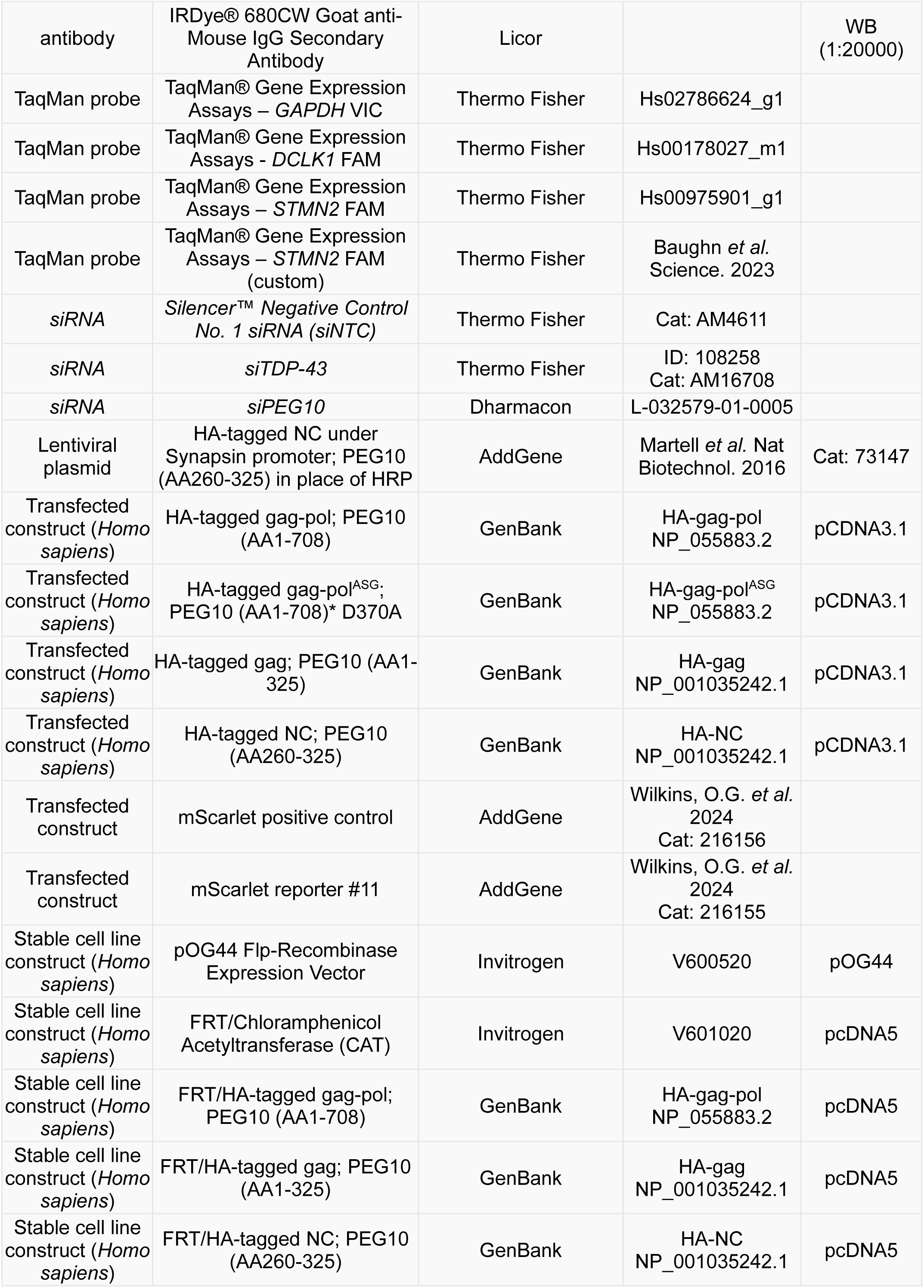
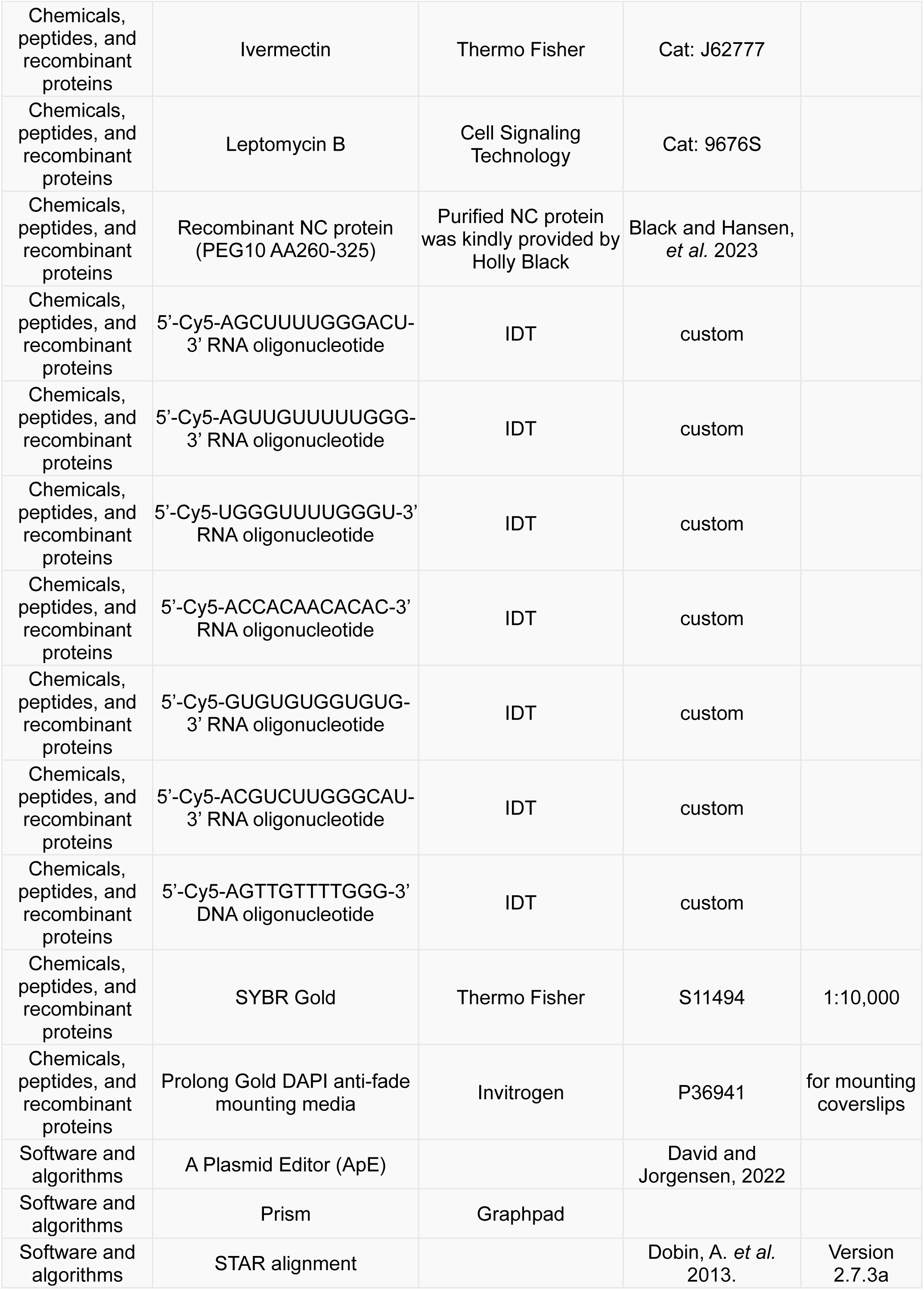
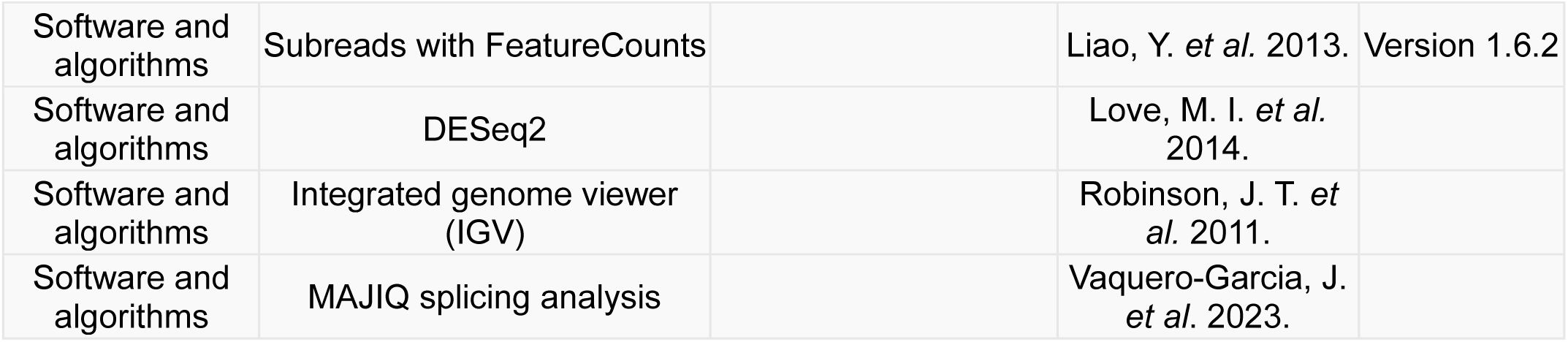

### Molecular Cloning

Constructs for stable cell line generation in Flp-In-293 cells and constructs for transient transfection were designed with a CMV promoter, and HA-NC construct for iNeuron expression was designed with a Synapsin promoter using Gibson or restriction/ligation cloning (see list of constructs). Plasmids were transformed into chemically competent DH5ɑ *E. coli* cells (Invitrogen). Transformed *E. coli* were plated on 100 μg/mL carbenicillin (Gold Biotechnology, cat: C-103–5) LB agar (Teknova, cat: L9115) plates overnight at 37 °C. Single colonies were grown overnight in 7 mL LB Broth (Alfa Aesar, cat: AAJ75854A1) with 1 x carbenicillin at 37 °C with shaking at 220 rpm. The following day, 5 mL of the shaking cultures were mini-prepped (Zymo, cat: D4212), and 1 mL was used to generate glycerol stocks of each plasmid. Plasmids were sent for sequencing via Azenta or Plasmidsaurus and were confirmed via alignment of idealized plasmids in APE^106^. To generate plasmid stocks, 50 mL shaking cultures were grown overnight from a glycerol stock or using 5 mL of overnight culture as previously described. The overnight culture was midi-prepped (Zymo, cat: D4200) to obtain a purified plasmid sample. Plasmid concentration was determined by nanodrop and were stored for transfections at 1 μg/μL in sterile water.

### Cell Culture

Mammalian cells were maintained in the Biochemistry Cell Culture Facility at CU Boulder (RRID:SCR_018988). HEK293 and SH-SY5Y cells were grown in complete media (DMEM (Gibco, cat: 12800082), 10% FBS (Atlas Biologicals, cat: F-05000-DR), GlutaminePlus (Bio-techne Sales Corporation cat: R90210), and Penicillin-Streptomycin (Invitrogen, cat: 15140163). Once cells reached 90-100% confluency, cells were washed with 1 x PBS then treated with Trypsin-EDTA (0.05%) (Invitrogen, 25300120), until cells began to detach. Complete media was added to harvest the cells. When plating for an experiment, live cells were counted via hemacytometer or using an automated cell counter (Corning, CytoSMART).

Human embryonic stem cells (hESCs, H9 strain) containing a stably integrated, doxycycline-inducible NGN2 cassette for neuron differentiation were grown as patches of cells on hESC-qualified Matrigel® Matrix (Corning, cat: 354277) in TeSR-E8 media (StemCell Technologies, cat: 05990) in 6-well plates. Once cell patches reached ∼70% confluency, cells were split for maintenance. Cells were washed with 1 mL of sterile 1 x PBS with 0.5 mM EDTA (Sigma-Aldrich, cat: E5134-500G) and another 1 mL was added and cells were kept at room temperature until cell patches began detaching. The 1 x PBS with 0.5 mM EDTA was carefully aspirated off and cell clumps were resuspended in TeSR-E8 media with a 5 mL pipette and distributed to new wells.

### Induced Neuron Differentiations

When splitting hESCs for neuron differentiation, cells were washed with 1 X PBS then incubated for 5 minutes with Accutase cell dissociation reagent (Gibco, cat: A1110501), until cells begin to detach. Cells were resuspended in ND1 media (DMEM (Gibco, cat: 12800082) and F12 (Gibco, cat: 21700075) at a 1:1 ratio, supplemented with 1 x N2 (Gibco, cat: 17502048), 1 x BDNF (VWR, cat: 10781-154), 1 x NT-3 (VWR, cat: 10781-168), 1 x MEM non-essential amino acids (Gibco, cat: 11140050), 0.2 𝜇g/mL Laminin (Sigma Aldrich, cat: L2020-1MG), and 8 𝜇g/mL Doxycycline (RPI, cat: D43020).

While still in suspension, cells were counted and then plated on Matrigel-coated coverslips in 12-well plates for IF staining or in Matrigel-coated 6-well plates without coverslips for western blot. Cells were plated in wells containing ND1 media supplemented with 10 μM ROCK inhibitor (Fisher Scientific, cat: BDB562822). After 24 hours, the media was replaced with ND1 media without ROCK inhibitor. 24 hours later, the media was replaced with ND2 media (Neurobasal media (Gibco, cat: 21103049) supplemented with B27 (Gibco 17504001), 1 X GlutaMAX supplement (Gibco, cat: 35050061), BDNF, NT-3, and 8 𝜇g/mL Doxycycline with 0.5 μg/mL puromycin (Thermo Fisher Scientific, cat: A1113803). 48 hours later, media was replaced with ND2 media without puromycin. 48 hours later, media was replaced with ND2 media. 24 hours later, cells were harvested for IF staining or for western blot.

### Cell Transfections

Cells were seeded in 12 or 6-well plates and grew until they reached ∼70% confluency. Cell media was changed to warmed transfection media containing DMEM, 10% FBS, and GlutaminePlus. Cells were transfected with 1 μg (12-well plate) or 2.5 μg (6-well plate) plasmid DNA in Lipofectamine 2000 (Invitrogen, cat: 11668027) and Opti-Mem medium (Invitrogen, cat: 11058021), according to manufacturer’s instructions. 48 hours after transfection, cells were harvested for subsequent experiments.

### Stable Cell Line Generation

Flp-In-293 cells (Invitrogen, cat: R75007) were cultured according to manufacturer instructions in complete media with zeocin (Gibco, cat: R25001). Cells were grown in a 6-well plate until they reached ∼70% confluency and zeocin media was removed and replaced with complete media. Cells were then transfected with 500 ng of pcDNA5/FRT plasmids or positive transfection control, pcDNA5/FRT chloramphenicol acetyltransferase (CAT) (Invitrogen, cat: V601020), in combination with 500 ng pOG44 (Invitrogen, cat: V600520) in Lipofectamine 2000 and Opti-Mem medium, according to manufacturer’s instructions. One well was also not transfected to act as a control for stable selection.

The following day, media was replaced with complete media. When cells reach 100% confluency in the 6-well plate, cells were transferred to a 10 cm dish. Once cells adhere to the 10 cm dish, media was replaced with complete media with 100 μg/mL hygromycin B (Gibco, cat: 10687010) to select for transfected cells. Cells were checked and media was replaced every 3-4 days until all untransfected cells had died. The stable cells were then split into a new 10 cm dish and continued to undergo hygromycin B selection for another week. Stable cell lines were plated in 12-well plate and harvested for western blot to assay for expression of our protein of interest. The HA-gag-pol stable cell line was further selected via single cell population. These cells were plated in a 96 well plate at ¼ cell per well to ensure some wells only had a single cell to start. These cells were grown up from that single cell until western blot could be performed to ensure stable cell line generation.

### Immunofluoresence

HEK293 and SH-SY5Y cells were plated onto Alcian blue-treated (Newcomer Supply, cat: 1002 A) round coverslips (Electron Microscopy Sciences, cat: 7223101SP) in complete media and transfected for 48 hours before harvesting, or were plated 48 hours post-transfection onto Alcian blue-treated coverslips in complete media with only 5% FBS and left to adhere overnight.

For NC trafficking, Flp-in-293 CAT or Flp-in-293 HA-NC-expressing cells were plated on coverslips and then incubated with either 25 μM ivermectin (Thermo Fisher, cat: J62777) or 50 nM Leptomycin B (CST, cat: 9676S) for 2 hours to inhibit trafficking pathways.

When ready to harvest, coverslips were washed with PBS then fixed with 100 μL of 4% PFA (Thermo Fisher Scientific, cat: 28906). Coverslips were then either incubated in 1% PFA for overnight storage or washed three times in 1 x PBS. Cells were permeabilized in 0.25% Triton-X (Sigma Aldrich, cat: X100) in PBS and incubated in blocking buffer containing 7.5% BSA (Gibco, cat: 15260037) diluted to 5% in PBS-T (0.1% Tween, (VWR, cat M147-1L)) for 30 min. Cells were incubated for 1 hour in primary antibody diluted in blocking buffer. Coverslips then underwent three 5 min washes in 1 x PBS-T.

Coverslips were incubated in secondary antibodies in blocking buffer for 1 hour in the dark. Cells underwent three more 5 min 1 x PBS-T washes and were then rinsed three times in DEPC water (Fisher Scientific, cat: AM9915G). Microscope slides (VWR, cat: 12-544-2) were cleaned with 70% ethanol and prepared for mounting. After sufficient drying, 5 μL of Prolong Gold DAPI anti-fade mounting media (Invitrogen, cat: P36941) was added to microscope slides. Coverslips were then mounted onto the microscope slides by flipping coverslips cell-side-down onto the mounting media and lightly pressing down to ensure the entire coverslip has been mounted. Microscope slides were then cured overnight in the dark at room temperature before imaging.

### Microscopy

Microscope slides were imaged on a Nikon A1R or a Nikon AXR laser scanning confocal microscope, through the Biofrontier Institute’s Advanced Light Microscopy Core at CU Boulder (RRID: SCR_018302). Imaging of SH-SY5Y cells and iNeurons for analyses was performed with a 20X air objective, and iNeurons for visualization with a 100X oil objective using NIS Elements software and all image panels were generated using Fiji/ImageJ^107^. For NRG3 analysis in SH-SY5Y cells, images were taken using Z-stacks with 0.5 μm step size for 11 total steps to generate whole cell images. All images for each individual experiment were taken at the same settings to ensure comparability between conditions of the same experiment. Contrast was enhanced in panel images using Fiji/ImageJ^107^ for ease of visualization, but this enhancement was not used for quantification of images. For all enhancements, the same settings were used across all images within the same experiment.

Flp-in-293 cell lines transfected with mScarlet reporters^74^ were imaged in their original cell culture dishes using a EVOS microscope with 20x objective. Cells were observed in transmitted light to ensure similar cell densities and mScarlet images were taken in the TexasRed channel.

### Analysis of Microscopy Images

For analysis of sub-cellular HA-NC localization in Flp-in-293 cells, at least 20 images of each condition were used for analysis using Matlab software. Nuclei were identified by DAPI intensity, and the cytoplasm was identified using tubulin intensity. The nuclear mask was removed from inside the cytoplasmic mask to generate a mask of just the cytoplasm. Regions of interest (ROI) were defined as the nuclear or cytoplasmic mask, then used to generate mean intensities of HA signal for each ROI as a ratio through Matlab.

For quantification of NRG3 and MAP2 intensity in transfected SH-SY5Y cells, images were analyzed on Fiji/ImageJ^107^. Z-stack images were compiled into a single image in which the maximum intensity was compiled for each channel per slice. The cytoplasm was identified using MAP2 signal intensity to generate manual ROIs of the entire cell. The NRG3 and MAP2 mean and maximum signal per cell was generated in Fiji/ImageJ^107^ by measuring the signal intensity of each cell/ROI.

Sholl analysis of iNeurons was performed using the Neuroanatomy/SNT plugin on Fiji/ImageJ^107^. iNeurons were skeletonized using the MAP2 channel to trace dendrites and Sholl analysis was performed with a radius step size of 10 micrometers. For each iNeuron skeletonized for Sholl analysis, each branch path was labeled as a primary, secondary, or tertiary dendrite depending on where it branched and those paths were used to measure the path length in micrometers of each dendrite.

Quantification of NRG3 intensity along dendrites was determined by generating lines along MAP2 intensity via the Neuroanatomy/SNT plugin on Fiji/ImageJ^107^ to trace dendrites. These dendritic lines were then used to generate ROIs that were measured at each point along the line to generate a line profile for both the NRG3 and MAP2 channels. At each point, the NRG3 over MAP2 signal intensity was quantified.

### Fluorescent Electrophoretic Mobility Shift Assays (fEMSAs)

Purified NC protein^23^ was thawed on ice, centrifuged for 15 minutes at 16,000 x g at 4C, and the supernatant was collected. The concentration of NC protein was quantified using BCA. For individual fEMSAs, NC protein underwent a serial dilutions to generate a range of 0 to 10μM of NC protein per reaction. NC protein was combined with 0.5 nM of 5’ Cy5-labeled RNA or DNA probe (IDT, see list of chemicals) per reaction and LICOR mobility shift assay binding buffers (LICORbio, cat: 829-07910) for a final concentration of 10 mM Tris, 50 mM KCl, 1 mM DTT (pH 7.5), 5 mM DTT, 0.5% Tween 20, 0.05% NP-40, 10 mM EDTA, pH 8, and 0.5 nM RNA probe per reaction. Reactions were incubated in the dark for 30 minutes. 6% Novex TBE gels (Invitrogen, cat: EC6265BOX) were pre-run without samples in 0.5 x TBE running buffer (VWR, cat: 0658-20L) for 15 min at 70V. After the samples finished incubating, LICOR orange loading dye (LICORbio, cat: 927-10100) was added to a final concentration of 1x in each sample and samples were loaded and run for 45 min at 100V in the dark at room temperature. Gels were visualized using a LICOR Odyssey CLx using the 700 nm channel. To generate images for publication, western blots were exported from ImageStudio as .png files and cropped in Adobe Illustrator. Uncropped blots are shown in Figure S6.

### Western Blotting

Cells were collected and centrifuged at 300 x g for 5 min to harvest cell pellets, which were then washed two times with PBS. After washing, cell pellets were lysed in urea lysis buffer (8 M urea (Fisher Chemical, cat: U15500), 75 mM NaCl (Honeywell Fluka, cat: 6003219), 50 mM HEPES (Millipore Sigma, cat: H3375) pH 8.5, 1 x tab cOmplete Mini EDTA-free protease inhibitor cocktail tablet (Roche, cat: 11836170001)). Lysed cells were briefly vortexed, centrifuged on a tabletop centrifuge to collect the liquid, and let sit at room temperature for 10 minutes, then kept overnight at −20°C. After thawing the next day, lysate was centrifuged for 10 min at 21,300 × g and the supernatant stored for western blot.

Protein was quantified by BCA (Pierce, cat: 23227), which was used to equalize protein input across lanes of western blots. Western blot samples were prepared with 1x Laemmli sample buffer supplemented with βME (Sigma Aldrich, cat: M3148), and urea lysis buffer before SDS-PAGE. Samples were run in NuPage MES Running Buffer (Invitrogen, cat: NP000202, diluted to 1x in DI water) on a 4 to 12% NuPage Bis-Tris gel (Invitrogen, cat: NP0321) at 120V for 60 min. Following SDS-PAGE, the gel was wet transferred onto nitrocellulose membrane (Amersham Protran, cat: 10600009) for 45 minutes at 15V using an Invitrogen mini-blot module.

After transfer, membranes were blocked using LICOR Intercept blocking buffer (LICORbio, cat: 927–70001) diluted 1:1 with TBST (50 mM Tris-HCl (MP Biomedicals, cat: MP04816116) pH 7.4, 150 mM NaCl, 0.1% Tween) for 1 hour at room temperature. Membranes were incubated in primary antibody overnight at 4 °C in 1x TBST. Membranes were washed in 1 x TBST 3 times in 5-min intervals. Membranes were then incubated in LICOR secondary antibody in 1 x TBST for 1 hour in the dark. Membranes were washed in 1x TBST 3 more times in 5-min intervals, then the nitrocellulose membranes were visualized using a LICOR Odyssey CLx. Data analysis was performed using LICOR ImageStudio Software. To generate images for publication, western blots were exported from ImageStudio as .png files and cropped in Adobe Illustrator. Uncropped blots are shown in Figure S6.

### RNA Isolation in SH-SY5Y Cells

SH-SY5Y cells were plated at 2.5*10^5^ cells per well of a 12-well plate. When cells reached 70% confluency, media was replaced with warmed transfection media. Cells were then transfected in duplicate wells with control *siNTC* (Thermo Scientific, cat: AM4611), *siTDP-43* (Thermo Scientific, cat: AM16708), *siPEG10* (Dharmacon, cat: L-032579-01-0005), or HA-NC plasmid in Lipofectamine 2000 with Opti-Mem medium, according to manufacturer’s instructions. Each well was harvested for western blot or RNA isolation 48 hours after transfection. Western blotting was performed to confirm expression of HA-NC or knockdown of TDP-43 or PEG10. For RNA isolation, cells were pelleted and RNA was extracted using the RNeasy Mini Kit (Qiagen, cat 74106) with on-column DNAse digestion. Extracted RNA was quantified using NanoDrop and stored at −80°C until samples were ready to prepare for sequencing.

### RNA-Sequencing and Analysis of SH-SY5Y Samples

RNA samples were diluted to 550 ng per 50 uL and sent for Poly-A Selected Total RNA Library paired-end sequencing at Anschutz Medical Campus, performed on an Illumina NovaSEQ X. Sequencing produced between 13,649 – 18,643 Mbases per sample with a mean quality ∼38.9 and 94.78 >=Q30. Quality of the reads was determined by FastQC version 0.11.8 before and after using Trimmomatic version 0.36^108^ to trim any leftover read ends, with 32 – 60 million reads left per sample after trimming. Reads were mapped to GRCh38.p13 using STAR version 2.7.3a^109^. All alignment files were indexed, sorted, and underwent feature count generation using Subread version 1.6.2^110^ including FeatureCounts.

Feature counts were imported into R Studio^111^ for further analysis and DESeq2^112^ analysis for differentially expressed genes. Genes with zero expression were filtered out and lfcShrink was used to apply fold change shrinkage. Normalized counts with an adjusted p-value of 0.05 were evaluated for top hits. Volcano plots of the top differentially expressed (DE) genes were generated, with a log_2_foldchange cutoff of 0.1 and an adjusted p-value of 0.05. Heatmaps of the top DE genes were generated by making a table with all significant results and pulling the top 25 DE genes across all conditions relative to *siNTC*. Using log_2_foldchange > 0.1 and log_2_foldchange < −0.1, heatmaps of the top 25 DE genes accumulated or depleted, respectively, were generated.

Pathway analysis was performed in R Studio using enrichGO^83^ with a log_2_foldchange cutoff of 0.1 and a p-value of 0.05 of significantly changed genes for each pairwise analysis. The top five pathways by p-value were visualized, as well as the top 5 accumulated or depleted pathways. Heatmaps were generated of the top 25 DE genes within GO-term enrichment pathways.

Splicing analysis was performed using MAJIQ^85,113^ and results were visualized using the MAJIQ Voila Viewer with a deltapsi threshold of 0.1 and confidence interval cutoff of 0.9. Splice variant classification analysis was performed using MAJIQ classifier. A venn diagram of splice changes specific to each condition relative to *siNTC* was generated by comparing all genes in MAJIQ Voila Viewer with a deltapsi threshold of 0.1 and confidence interval cutoff of 0.9. Further validation of splice changes was performed by visualizing splice variants in IGV^114,115^. To generate images for publication, IGV screenshots were exported and cropped in Adobe Illustrator. Uncropped blots are shown in Figure S7.

### RT-PCR and qPCR

For RT-PCR and qPCR, isolated RNA from SH-SY5Y cells was reverse transcribed into cDNA using a High-Capacity cDNA Reverse Transcription Kit (Applied Biosystems, cat: 43-688-14), according to manufacturer instructions. cDNA was quantified via NanoDrop and diluted to 500 ng/μL for RT-PCR or 10ng/μL for qPCR. For RT-PCR, all reactions were performed using 2X Phusion Master Mix (Thermo Fisher, cat: F531L), forward and reverse primers^58^, and 1 μg cDNA per reaction, diluted in water. Final PCR products were combined with 6 X loading dye (Thermo Fisher, cat: R0631) and analyzed by electrophoresis on 2% agarose gels for 20 minutes at 125V. Gels were incubated with 1 X SYBR Gold for 30 minutes and imaged on an Azure Biosystems gel imager.

For each biological replicate, qPCR was performed in triplicate. PerfeCTa qPCR FastMix II low ROX master mix (Quantabio, cat: 95120-012) and TaqMan probes (VIC-labeled GAPDH control and FAM-labeled probe of interest) were thawed on ice, then combined with 10 ng of cDNA per reaction according to manufacturer’s instructions, then added to each well of a MicroAmp Fast 96-well reaction plate (Applied Biosystems, cat: 4346907). The plate was sealed with an optical adhesive cover (Applied Biosystems, cat: 4360954), centrifuged for 30 seconds at 100 x g, and the reactions were run on a QuantStudio 6 machine according to PerfeCTa Master Mix guidelines. To analyze results, Ct values were obtained for GAPDH and each probe of interest. To determine the relative expression of *STMN2* long or short isoforms, the 2^-ΔΔCt^ value was calculated for each condition. Technical replicates that had Ct values that were undetermined were not included in the final analysis.

### ALS Patient Sample Gene Expression Analysis

RNA-Seq reads from post-mortem lumbar spinal cord samples from the Target ALS: New York Genome Center dataset were obtained, aligned, and initially analyzed according to previous publication^23^. In brief, samples included 126 ALS cases, which include cases with identified genetic mutation, and 16 non-neurological control (NNC) post-mortem controls, as well as one *UBQLN2*-mediated case of ALS-FTD. BAM files and DESeq2 previously published^23^ were used to query specific transcripts for their abundance and splicing profile in IGV.

## Supplemental Figures

**Figure S1.**
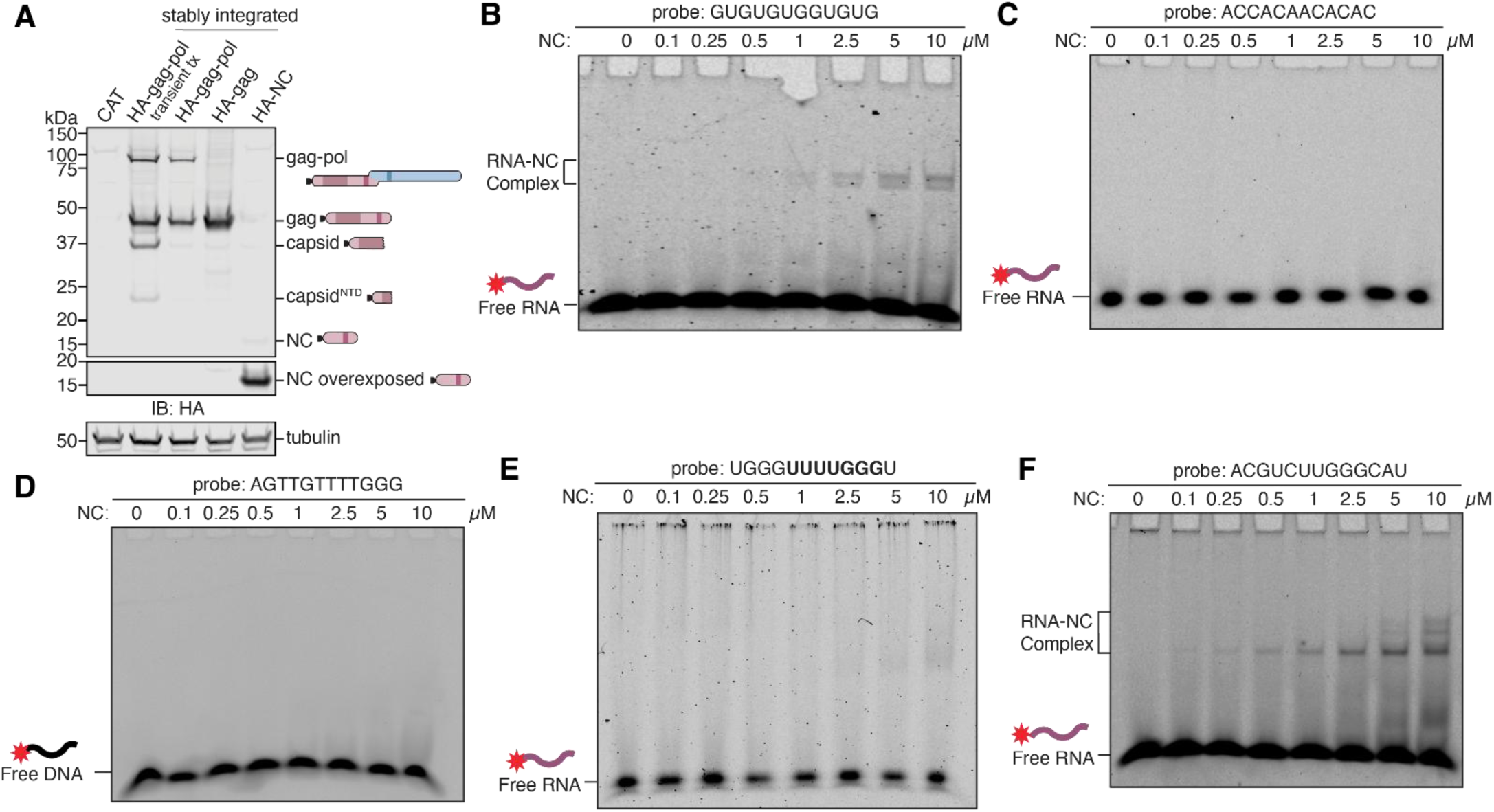
Stable cell line generation and additional fEMSAs showing NC-specific RNA-binding preference, related to Figure 1D-F. **(A)** Western blot of whole cell lysate from Flp-In-293 cells stably expressing control chloramphenicol acetyltransferase (CAT), HA-gag-pol, HA-gag, or HA-NC. In lane 2, whole cell lysate of Flp-In-293 cells transiently transfected with HA-gag-pol for 24 hours was performed to compare PEG10 expression and self-processing levels before and after stable integration. Lysate was probed by western blot for HA and tubulin. HA-tagged NC runs at about 15 kDa and was overexposed for ease of visualization at bottom. **(B)** fEMSA of PEG10 NC incubated with 0.5 nM GUGUGUGGUGUG RNA oligo with a 5’ Cy5.5 probe. (n=3). **(C)** fEMSA of PEG10 NC incubated with 0.5 nM ACCACAACACAC RNA oligo with a 5’ Cy5.5 probe. (n=3). **(D)** fEMSA of PEG10 NC incubated with 0.5 nM AGTTGTTTTGGG DNA oligo with a 5’ Cy5.5 probe. (n=2). **(E)** fEMSA of PEG10 NC incubated with 0.5 nM UGGGGUUUUGGGU RNA oligo with a 5’ Cy5.5 probe. (n=2). **(F)** fEMSA of PEG10 NC incubated with 0.5 nM ACGUCUUGGGCAU RNA oligo with a 5’ Cy5.5 probe. (n=2).

**Figure S2.**
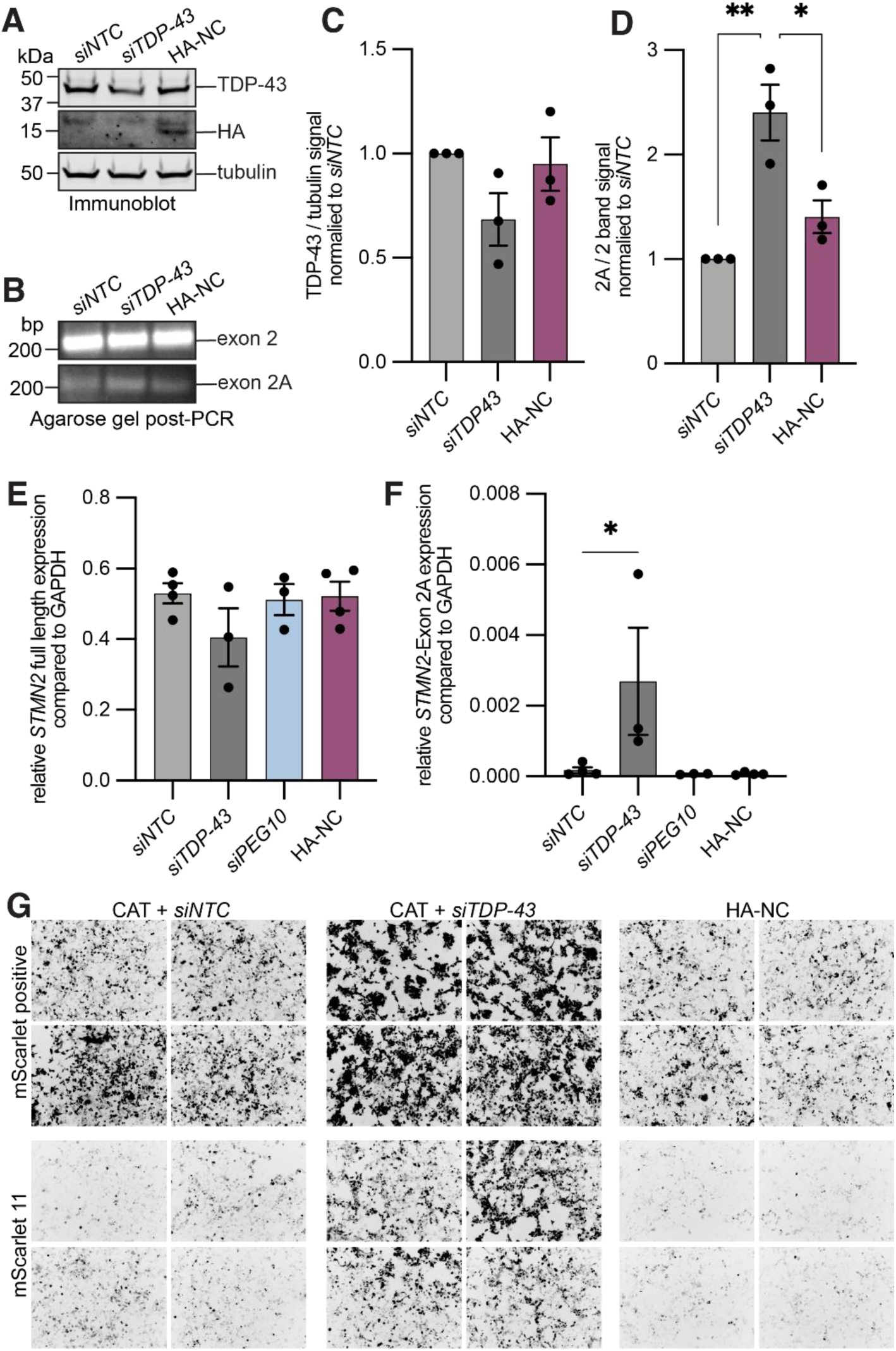
NC does not regulate splicing of *STMN2* in the same manner as TDP-43, related to Figures 2 and 4. **(A)** Representative western blot of SH-SY5Y cells transfected with *siNTC, siTDP-43*, or HA-NC. Cells were harvested 48 hours post transfection and probed for HA, TDP-43, and tubulin. (n=3). **(B)** Representative PCR results of SH-SY5Y cells transfected as in (A) for *STMN2* exon 2A or 2 visualized by 2% agarose gel stained with SYBR Gold (n=3). **(C)** Quantification of western blot results from (A). (n=4). There are no significant differences in TDP43 levels. **(D)** Quantification of agarose gel comparing the presence of *STMN2* exon 2A to exon 2. For (C-D), a one-way ANOVA was performed and compared to the means of each other. **(E)** qPCR for full-length *STMN2,* **(F)** qPCR for *STMN2* exon 2A. For (E-F), a one-way ANOVA was performed and compared to the mean of si*NTC*. (n=3 or 4). **(G)** Representative images of control CAT or HA-NC stable cell lines transfected with mScarlet reporters of TDP-43-dependent cryptic exon inclusion. mScarlet positive control, and reporter #11 were selected for testing. CAT HEK293 cells were transfected with either *siNTC* or *siTDP-43* for 24 hours, then all cell lines were transfected with mScarlet splice reporter constructs for an additional 24 hours. Images were taken at 20X magnification on an EVOS microscope, with 4 images per condition to show representative fields of view (n=3). *p<0.05, **p<0.01, ***p<0.001, and ****p<0.0001.

**Figure S3.**
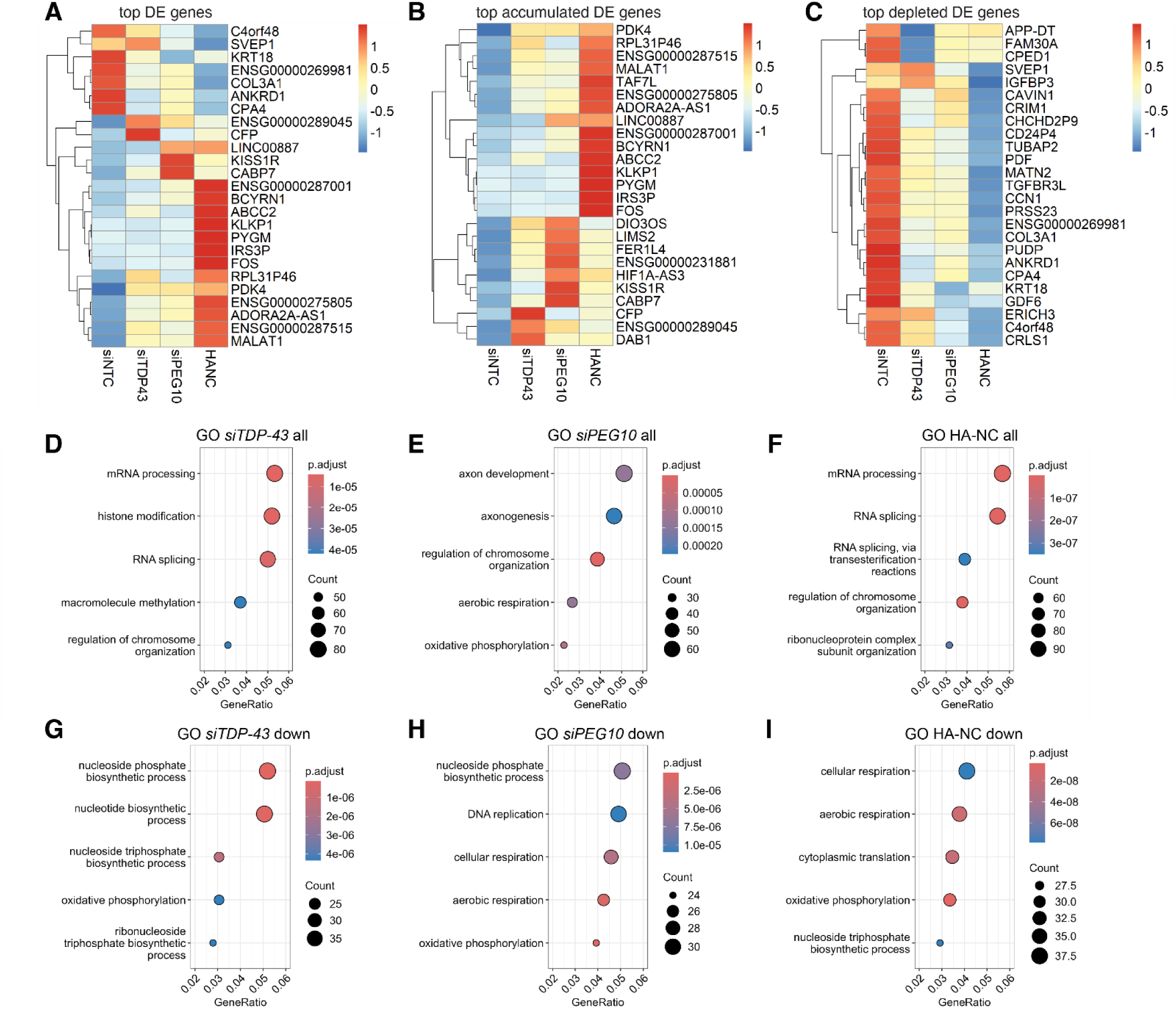
NC leads to widespread changes to gene regulation pathways, related to Figure 3. **(A-C)** Heatmap of **(A)** the most differentially expressed (DE) genes across all conditions **(B)** most accumulated DE genes across all conditions and **(C)** most depleted DE genes across all conditions. (n=4). For each heatmap, the top genes across all conditions relative to *siNTC* were combined. Genes were then ranked according to log_2_FoldChange across the combined dataset. Genes were then color coded by row Z-score. **(D-F)** Top changed pathways of gene expression changes in either direction in cells transfected with **(D)** *siTDP-43,* **(E)** *siPEG10*, or **(F)** HA-NC by GO-term enrichment analysis. **(G-I)** Top downregulated pathways of gene expression changes in cells transfected with **(G)** *siTDP-43,* **(H)** *siPEG10*, or **(I)** HA-NC by GO-term enrichment analysis. The top five GO-terms using a log_2_FoldChange cutoff of 0.1 were ranked by adjusted p-value. Adjusted p-value is shown by color, and the size of the datapoint reflects the number of genes enriched in the pathway.

**Figure S4.**
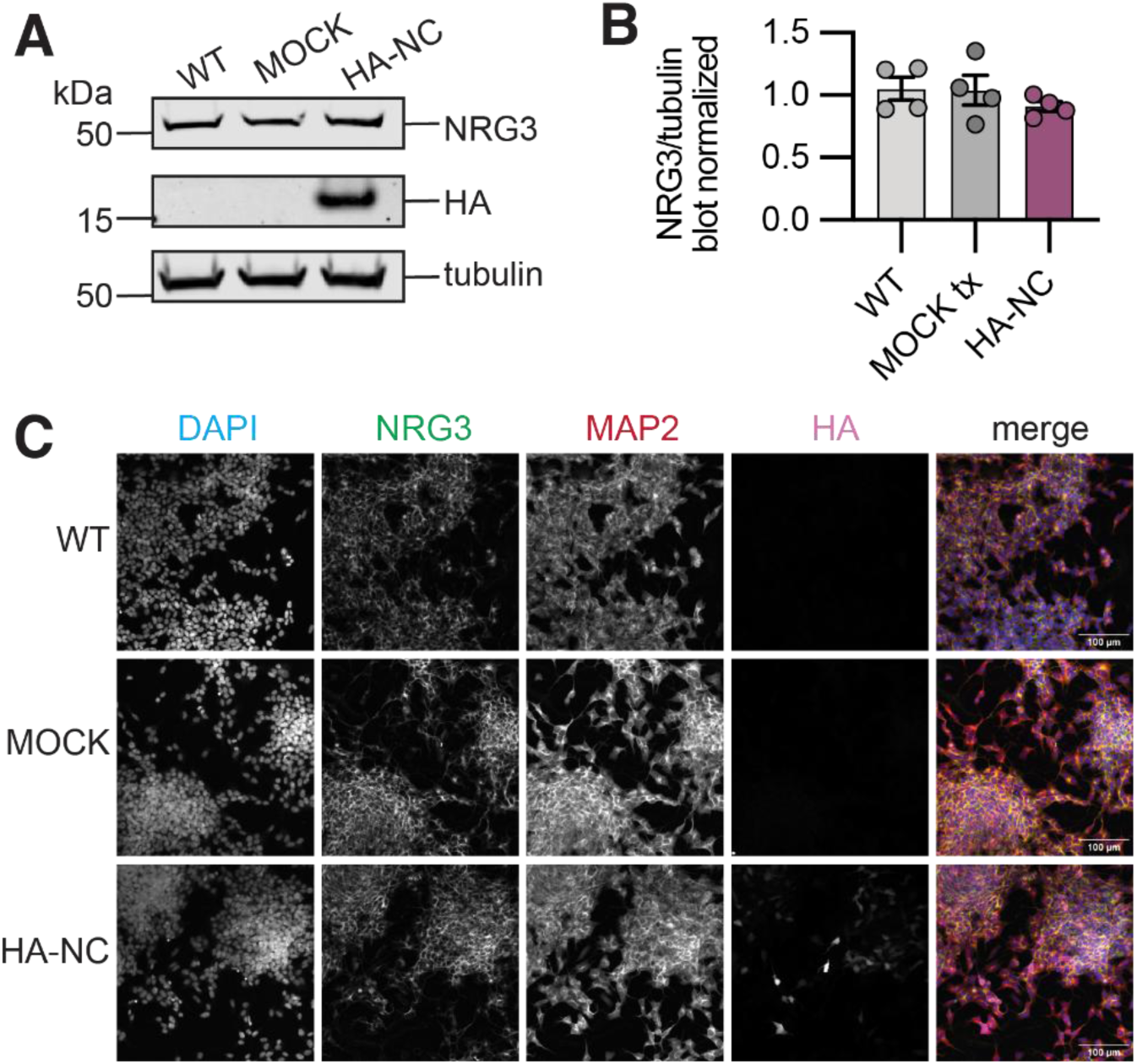
NRG3 abundance is not altered in bulk lysate of NC-expressing SH-SY5Y cells, related to Figure 5. **(A)** Representative western blot of WT SH-SY5Y cells compared to cells that have been mock transfected with lipofectamine alone (MOCK), or transfected with HA-NC.Cells were harvested 48 hours post transfection and probed for HA, NRG3, and tubulin. (n=4). **(B)** Quantification of western blot results from A. (n=4). There are no significant differences in NRG3 levels by one-way ANOVA. **(C)** Representative images from IF staining of WT SH-SY5Y cells compared to MOCK or HA-NC transfected cells. Cells were stained with NRG3, MAP2, and HA and coverslips were mounted with Prolong Gold with DAPI. Z-stack images were taken on a Nikon AXR at a 2048 size at Nyquist. Scale bar = 10 μm, and contrast has been increased for ease of visualization. (n=3).

**Figure S5.**
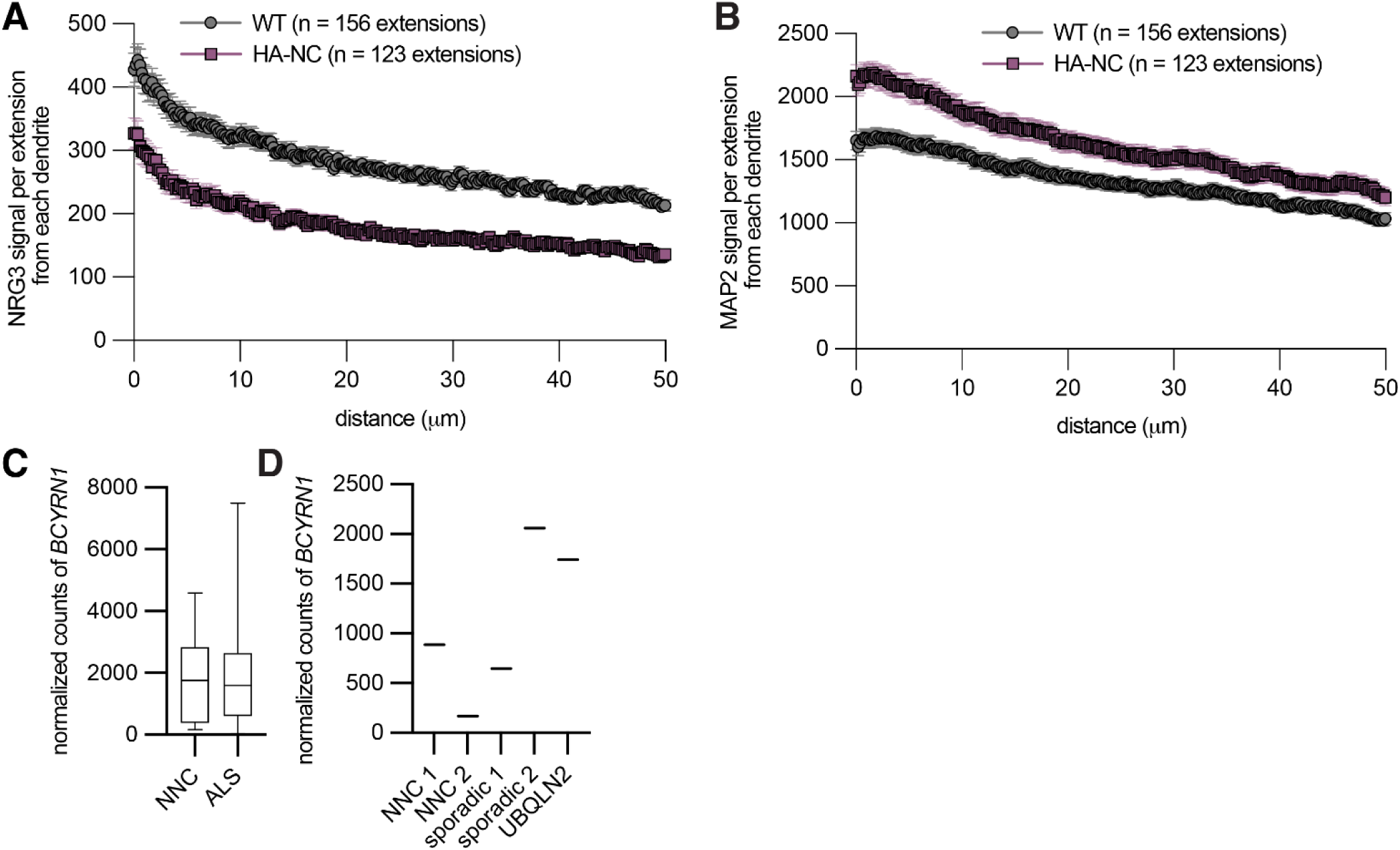
MAP2 is not depleted in iNeurons expressing NC and BCYRN1 is altered in human ALS, related to Figures 6 and 7. **(A-B)** mean **(A)** NRG3 or **(B)** MAP2 signal from dendrites starting at the branch point from the soma in differentiated iNeurons of either WT or HA-NC expression (n=1 quantified experiment). The number of extensions is shown in each legend. (C) Normalized counts of *BCYRN1* for NNC and sporadic ALS samples for Target ALS patient samples analyzed in Figure 7D-F. Counts were not significantly different by Student’s T test. (D) Normalized counts of *BCYRN1* for targetALS patient samples ananlyzed in Figure 7H, illustrating a similar expression profile with the exclusion of exon 9 in *UBQLN2*-mediated fALS and one sALS case. Datapoints are shown as lines to highlight that only one sample is evaluated per column.

**Figure S6.**
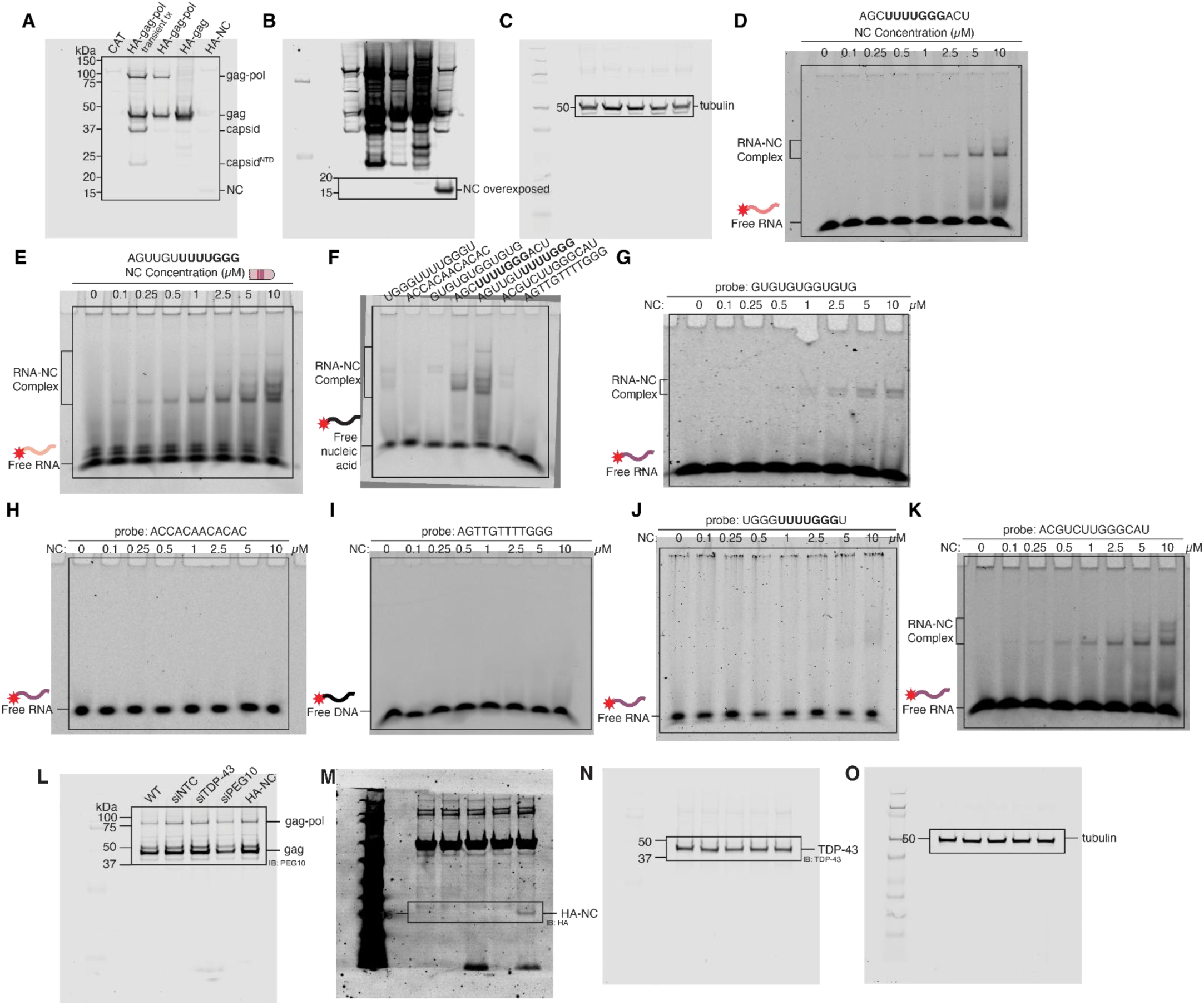
Uncropped western blots and fEMSAs, related to Figures 1 and 2. **(A-C)** Uncropped western blot from Figure S1A, **(D-K)** Uncropped fEMSAs from Figure 1E-G and Figure S1B-F, **(L-O)** Uncropped western blots from Figure 2A.

**Figure S7.**
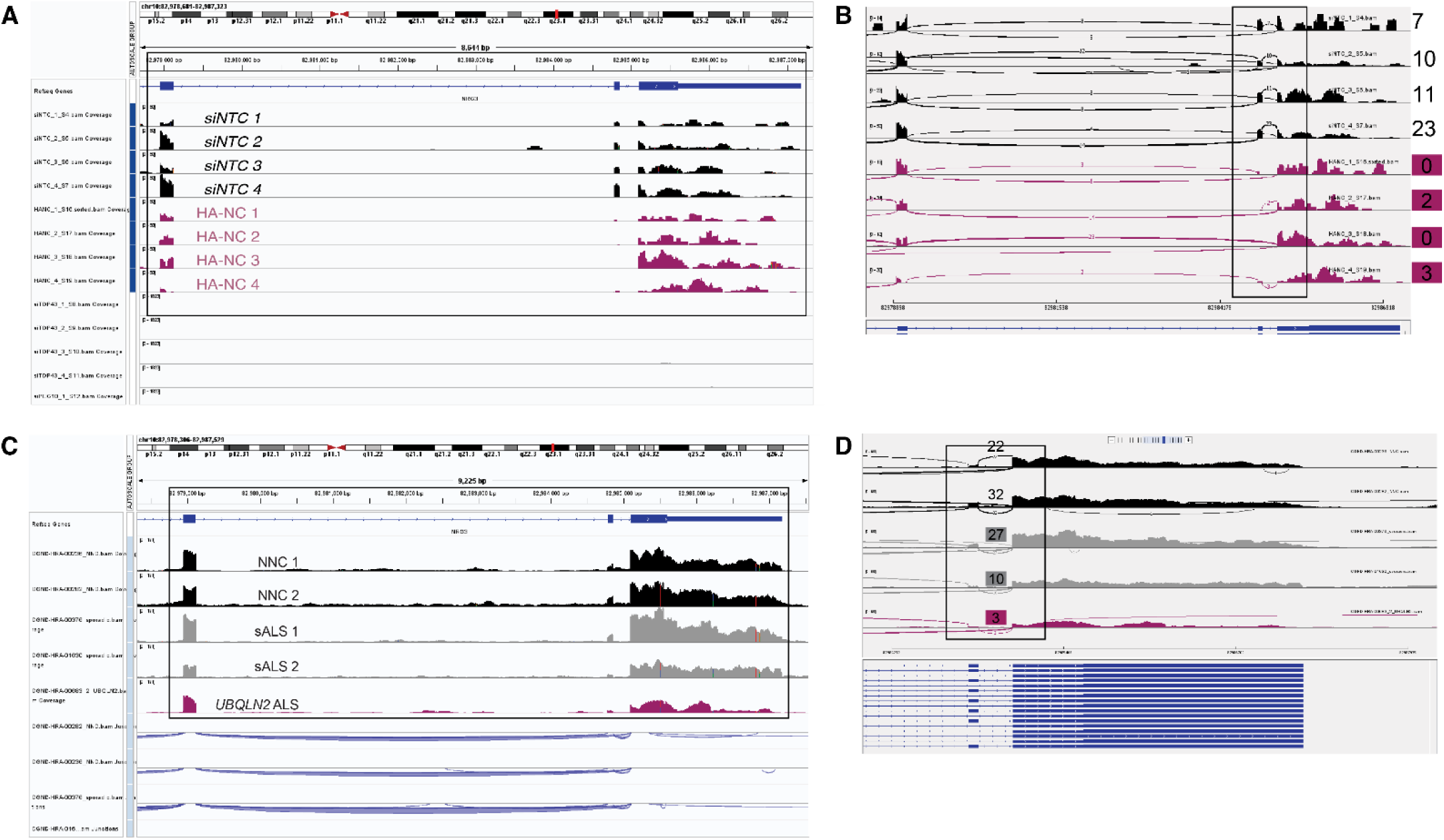
Uncropped IGV snapshots, related to Figures 4 and 7. **(A-B)** uncropped IGV snapshots from Figure 4F-G. **(C-D)** Uncropped IGV snapshots from Figure 7F.

**Table S1: Gene expression changes and pathway analysis of SH-SY5Y RNA-Seq data. (A-C)** Differentially expressed genes using a log_2_FoldChange cutoff of 0.1 and p adjusted value cutoff of 0.05 for all three relevant comparisons. **(D-F)** GO-term enrichments for NC-expressing SH-SY5Y cells compared to *siNTC*-expressing cells including upregulated genes, all significantly changed genes, or downregulated genes with a log_2_FoldChange cutoff of 0.1 and p adjusted value cutoff of 0.05. **(G-I)** GO-term enrichments for *siPEG10*-expressing SH-SY5Y cells compared to *siNTC*-expressing cells including upregulated genes, all significantly changed genes, or downregulated genes with a log2fold cutoff of 0.1 and p adjusted value cutoff of 0.05. **(J-L)** GO-term enrichments for *siTDP-43*-expressing SH-SY5Y cells compared to *siNTC*-expressing cells including upregulated genes, all significantly changed genes, or downregulated genes with a log_2_FoldChange cutoff of 0.1 and p adjusted value cutoff of 0.05.

**Table S2: Top splice changes between TDP-43 depletion, PEG10 depletion, and NC overexpression.** Top local splice variants (LSVs) for *siTDP-43*, *siPEG10*, and HA-NC transfected cells compared to siNTC using MAJIQ Splice changes were determined using a deltapsi cutoff of 0.1 and a confidence interval cutoff of 0.9 and all LSVs were compiled for each condition.

## References

1. Ravel-Godreuil, C., Znaidi, R., Bonnifet, T., Joshi, R. L. & Fuchs, J. Transposable elements as new players in neurodegenerative diseases. FEBS Letters 595, 2733–2755 (2021).

2. Whiteley, A. M. & Shepherd, J. D. Retrotransposons unplugged: Rewiring the nervous system and wreaking havoc. Neuron 114, 33–45 (2026).

3. Frost, B. & Dubnau, J. The Role of Retrotransposons and Endogenous Retroviruses in Age-Dependent Neurodegenerative Disorders. Annual Review of Neuroscience 47, 123–143 (2024).

4. Gorbunova, V. et al. The role of retrotransposable elements in aging and age-associated diseases. Nature 596, 43–53 (2021).

5. Brown, R. H. & Al-Chalabi, A. Amyotrophic Lateral Sclerosis. New England Journal of Medicine 377, 162–172 (2017).

6. Boillée, S., Velde, C. V. & Cleveland, D. W. ALS: A Disease of Motor Neurons and Their Nonneuronal Neighbors. Neuron 52, 39–59 (2006).

7. Ferrari, R., Kapogiannis, D., Huey, E. D. & Momeni, P. FTD and ALS: a tale of two diseases. Curr Alzheimer Res 8, 273–294 (2011).

8. Ling, S.-C., Polymenidou, M. & Cleveland, D. W. Converging Mechanisms in ALS and FTD: Disrupted RNA and Protein Homeostasis. Neuron 79, 416–438 (2013).

9. Ghasemi, M. & Brown, R. H. Genetics of Amyotrophic Lateral Sclerosis. Cold Spring Harb Perspect Med 8, a024125 (2018).

10. Ciechanover, A. & Kwon, Y. T. Degradation of misfolded proteins in neurodegenerative diseases: therapeutic targets and strategies. Exp Mol Med 47, e147–e147 (2015).

11. Ross, C. A. & Poirier, M. A. Protein aggregation and neurodegenerative disease. Nat Med 10, S10–S17 (2004).

12. Rubinsztein, D. C. The roles of intracellular protein-degradation pathways in neurodegeneration. Nature 443, 780–786 (2006).

13. Taylor, J. P., Brown, R. H. & Cleveland, D. W. Decoding ALS: from genes to mechanism. Nature 539, 197–206 (2016).

14. Genge, A., Wainwright, S. & Vande Velde, C. Amyotrophic lateral sclerosis: exploring pathophysiology in the context of treatment. Amyotrophic Lateral Sclerosis and Frontotemporal Degeneration 25, 225–236 (2024).

15. Tam, O. H. et al. Postmortem Cortex Samples Identify Distinct Molecular Subtypes of ALS: Retrotransposon Activation, Oxidative Stress, and Activated Glia. Cell Reports 29, 1164–1177.e5 (2019).

16. O’Neill, K. et al. ALS molecular subtypes are a combination of cellular and pathological features learned by deep multiomics classifiers. Cell Rep 44, 115402 (2025).

17. Bolger, I. et al. TDP-43 dysfunction leads to the accumulation of cryptic transposable element-derived exons, crypTEs, in iPSC derived neurons and ALS/FTD patient tissues. 2026.01.09.698641 Preprint at 10.64898/2026.01.09.698641 (2026).

18. Hou, Y. et al. TDP-43 chronic deficiency leads to dysregulation of transposable elements and gene expression by affecting R-loop and 5hmC crosstalk. Cell Reports 43, 113662 (2024).

19. Li, T. D. et al. TDP-43 safeguards the embryo genome from L1 retrotransposition. Science Advances 8, eabq3806 (2022).

20. Chang, Y.-H. & Dubnau, J. Endogenous retroviruses and TDP-43 proteinopathy form a sustaining feedback driving intercellular spread of Drosophila neurodegeneration. Nat Commun 14, 966 (2023).

21. Li, W., Jin, Y., Prazak, L., Hammell, M. & Dubnau, J. Transposable Elements in TDP-43-Mediated Neurodegenerative Disorders. PLoS One 7, e44099 (2012).

22. Whiteley, A. M. et al. Global proteomics of Ubqln2-based murine models of ALS. J Biol Chem 296, 100153 (2021).

23. Black, H. H. et al. UBQLN2 restrains the domesticated retrotransposon PEG10 to maintain neuronal health in ALS. eLife 12, e79452 (2023).

24. Pandya, N. J. et al. Secreted retrovirus-like GAG-domain-containing protein PEG10 is regulated by UBE3A and is involved in Angelman syndrome pathophysiology. Cell Reports Medicine 2, 100360 (2021).

25. Deng, H.-X. et al. Mutations in UBQLN2 cause dominant X-linked juvenile and adult-onset ALS and ALS/dementia. Nature 477, 211–215 (2011).

26. Mohan, H. M. et al. Endogenous retrovirus-like proteins recruit UBQLN2 to stress granules and shape their functional biology. Science Advances 11, eadu6354 (2025).

27. Mohan, H. M. et al. RTL8 promotes nuclear localization of UBQLN2 to subnuclear compartments associated with protein quality control. Cell. Mol. Life Sci. 79, 176 (2022).

28. Thumbadoo, K. M. et al. Hippocampal aggregation signatures of pathogenic UBQLN2 in amyotrophic lateral sclerosis and frontotemporal dementia. Brain 147, 3547–3561 (2024).

29. Williams, K. L. et al. *UBQLN2*/ubiquilin 2 mutation and pathology in familial amyotrophic lateral sclerosis. Neurobiology of Aging 33, 2527.e3–2527.e10 (2012).

30. Renaud, L., Picher-Martel, V., Codron, P. & Julien, J.-P. Key role of UBQLN2 in pathogenesis of amyotrophic lateral sclerosis and frontotemporal dementia. Acta Neuropathologica Communications 7, 103 (2019).

31. Matthews, A. M. & Whiteley, A. M. UBQLN2 in neurodegenerative disease: mechanistic insights and emerging therapeutic potential. Biochem Soc Trans 53, 823–833 (2025).

32. Brandt, J. et al. Transposable elements as a source of genetic innovation: expression and evolution of a family of retrotransposon-derived neogenes in mammals. Gene 345, 101–111 (2005).

33. Henriques, W. S. et al. The Diverse Evolutionary Histories of Domesticated Metaviral Capsid Genes in Mammals. Mol Biol Evol 41, msae061 (2024).

34. Campillos, M., Doerks, T., Shah, P. K. & Bork, P. Computational characterization of multiple Gag-like human proteins. Trends in Genetics 22, 585–589 (2006).

35. Kokošar, J. & Kordiš, D. Genesis and Regulatory Wiring of Retroelement-Derived Domesticated Genes: A Phylogenomic Perspective. Mol Biol Evol 30, 1015–1031 (2013).

36. Volff, J.-N. Cellular Genes Derived from Gypsy/Ty3 Retrotransposons in Mammalian Genomes. Annals of the New York Academy of Sciences 1178 233–243 (2009).

37. Clark, M. B. et al. Mammalian gene PEG10 expresses two reading frames by high efficiency −1 frameshifting in embryonic-associated tissues. J Biol Chem 282, 37359–37369 (2007).

38. Lux, A. et al. Human Retroviral gag- and gag-pol-like Proteins Interact with the Transforming Growth Factor-β Receptor Activin Receptor-like Kinase 1 * [boxs]. Journal of Biological Chemistry 280, 8482–8493 (2005).

39. Golda, M., Mótyán, J. A., Mahdi, M. & Tőzsér, J. Functional Study of the Retrotransposon-Derived Human PEG10 Protease. Int J Mol Sci 21, 2424 (2020).

40. Ayala, Y. M. et al. Structural determinants of the cellular localization and shuttling of TDP-43. J Cell Sci 121, 3778–3785 (2008).

41. Suk, T. R. & Rousseaux, M. W. C. The role of TDP-43 mislocalization in amyotrophic lateral sclerosis. Molecular Neurodegeneration 15, 45 (2020).

42. Winton, M. J. et al. Disturbance of Nuclear and Cytoplasmic TAR DNA-binding Protein (TDP-43) Induces Disease-like Redistribution, Sequestration, and Aggregate Formation. J Biol Chem 283, 13302–13309 (2008).

43. Colombrita, C. et al. TDP-43 and FUS RNA-binding Proteins Bind Distinct Sets of Cytoplasmic Messenger RNAs and Differently Regulate Their Post-transcriptional Fate in Motoneuron-like Cells *. Journal of Biological Chemistry 287, 15635–15647 (2012).

44. Ederle, H. & Dormann, D. TDP-43 and FUS en route from the nucleus to the cytoplasm. FEBS Lett 591, 1489–1507 (2017).

45. Luisier, R. et al. Intron retention and nuclear loss of SFPQ are molecular hallmarks of ALS. Nat Commun 9, 2010 (2018).

46. Piñol-Roma, S. & Dreyfuss, G. Shuttling of pre-mRNA binding proteins between nucleus and cytoplasm. Nature 355, 730–732 (1992).

47. Nakielny, S. & Dreyfuss, G. Transport of Proteins and RNAs in and out of the Nucleus. Cell 99, 677–690 (1999).

48. Wagstaff, K. M., Sivakumaran, H., Heaton, S. M., Harrich, D. & Jans, D. A. Ivermectin is a specific inhibitor of importin α/β-mediated nuclear import able to inhibit replication of HIV-1 and dengue virus. Biochem J 443, 851–856 (2012).

49. Wolff, B., Sanglier, J. J. & Wang, Y. Leptomycin B is an inhibitor of nuclear export: inhibition of nucleo-cytoplasmic translocation of the human immunodeficiency virus type 1 (HIV-1) Rev protein and Rev-dependent mRNA. Chem Biol 4, 139–147 (1997).

50. Wang, Y. et al. The distinct roles of zinc finger CCHC-type (ZCCHC) superfamily proteins in the regulation of RNA metabolism. RNA Biol 18, 2107–2126 (2021).

51. Summers, M. F. Zinc finger motif for single-stranded nucleic acids? investigations by nuclear magnetic resonance. Journal of Cellular Biochemistry 45, 41–48 (1991).

52. Steplewski, A. et al. MyEF-3, a Developmentally Controlled Brain-Derived Nuclear Protein Which Specifically Interacts with Myelin Basic Protein Proximal Regulatory Sequences. Biochemical and Biophysical Research Communications 243, 295–301 (1998).

53. Abed, M. et al. The Gag protein PEG10 binds to RNA and regulates trophoblast stem cell lineage specification. PLOS ONE 14, e0214110 (2019).

54. Segel, M. et al. Mammalian retrovirus-like protein PEG10 packages its own mRNA and can be pseudotyped for mRNA delivery. Science 373, 882–889 (2021).

55. Ray, D. et al. RNA-binding proteins that lack canonical RNA-binding domains are rarely sequence-specific. Sci Rep 13, 5238 (2023).

56. Arnold, E. S. et al. ALS-linked TDP-43 mutations produce aberrant RNA splicing and adult-onset motor neuron disease without aggregation or loss of nuclear TDP-43. Proceedings of the National Academy of Sciences 110, E736–E745 (2013).

57. Tollervey, J. R. et al. Characterizing the RNA targets and position-dependent splicing regulation by TDP-43. Nat Neurosci 14, 452–458 (2011).

58. Baughn, M. W. et al. Mechanism of STMN2 cryptic splice-polyadenylation and its correction for TDP-43 proteinopathies. Science 379, 1140–1149 (2023).

59. Ling, J. P., Pletnikova, O., Troncoso, J. C. & Wong, P. C. TDP-43 repression of nonconserved cryptic exons is compromised in ALS-FTD. Science 349, 650–655 (2015).

60. Melamed, Z. et al. Premature polyadenylation-mediated loss of stathmin-2 is a hallmark of TDP-43-dependent neurodegeneration. Nat Neurosci 22, 180–190 (2019).

61. Bryce-Smith, S. et al. TDP-43 loss induces cryptic polyadenylation in ALS/FTD. Nat Neurosci 28, 2190–2200 (2025).

62. Arnold, F. J. et al. TDP-43 dysregulation of polyadenylation site selection is a defining feature of RNA misprocessing in amyotrophic lateral sclerosis and frontotemporal dementia. J Clin Invest 135, (2025).

63. Zeng, Y. et al. TDP-43 nuclear loss in FTD/ALS causes widespread alternative polyadenylation changes. Nat Neurosci 28, 2180–2189 (2025).

64. Buratti, E. & Baralle, F. E. The multiple roles of TDP-43 in pre-mRNA processing and gene expression regulation. RNA Biology 7, 420–429 (2010).

65. Alami, N. H. et al. Axonal Transport of TDP-43 mRNA Granules Is Impaired by ALS-Causing Mutations. Neuron 81, 536–543 (2014).

66. Chu, J.-F., Majumder, P., Chatterjee, B., Huang, S.-L. & Shen, C.-K. J. TDP-43 Regulates Coupled Dendritic mRNA Transport-Translation Processes in Co-operation with FMRP and Staufen1. Cell Reports 29, 3118–3133.e6 (2019).

67. Kabashi, E. et al. Gain and loss of function of ALS-related mutations of TARDBP (TDP-43) cause motor deficits in vivo. Hum Mol Genet 19, 671–683 (2010).

68. Moens, T. G. et al. Amyotrophic lateral sclerosis caused by *FUS* mutations: advances with broad implications. The Lancet Neurology 24, 166–178 (2025).

69. Vance, C. et al. Mutations in FUS, an RNA processing protein, cause familial amyotrophic lateral sclerosis type 6. Science 323, 1208–1211 (2009).

70. Kim, H. J. et al. Mutations in prion-like domains in hnRNPA2B1 and hnRNPA1 cause multisystem proteinopathy and ALS. Nature 495, 467–473 (2013).

71. Zhao, M., Kim, J. R., van Bruggen, R. & Park, J. RNA-Binding Proteins in Amyotrophic Lateral Sclerosis. Mol Cells 41, 818–829 (2018).

72. Kapeli, K., Martinez, F. J. & Yeo, G. W. Genetic mutations in RNA-binding proteins and their roles in ALS. Hum Genet 136, 1193–1214 (2017).

73. Wood, H. TDP43 blocks misprocessing of STMN2 RNA. Nat Rev Neurol 19, 326–326 (2023).

74. Wilkins, O. G. et al. Creation of de novo cryptic splicing for ALS and FTD precision medicine. Science 386, 61–69 (2024).

75. Shaw, G., Morse, S., Ararat, M. & Graham, F. L. Preferential transformation of human neuronal cells by human adenoviruses and the origin of HEK 293 cells. The FASEB Journal 16, 869–871 (2002).

76. Prpar Mihevc, S., Baralle, M., Buratti, E. & Rogelj, B. TDP-43 aggregation mirrors TDP-43 knockdown, affecting the expression levels of a common set of proteins. Sci Rep 6, 33996 (2016).

77. Herzog, J. J., Deshpande, M., Shapiro, L., Rodal, A. A. & Paradis, S. TDP-43 misexpression causes defects in dendritic growth. Sci Rep 7, 15656 (2017).

78. Neeves, J. et al. An alternative cytoplasmic SFPQ isoform with reduced phase separation potential is up-regulated in ALS. Science Advances 11, eadt4814 (2025).

79. Widagdo, J. et al. Familial ALS-associated SFPQ variants promote the formation of SFPQ cytoplasmic aggregates in primary neurons. Open Biol 12, 220187 (2022).

80. Akiyama, T. et al. Aberrant axon branching via Fos-B dysregulation in FUS-ALS motor neurons. eBioMedicine 45, 362–378 (2019).

81. Mus, E., Hof, P. R. & Tiedge, H. Dendritic BC200 RNA in aging and in Alzheimer’s disease. Proceedings of the National Academy of Sciences 104, 10679–10684 (2007).

82. Zhao, J. et al. APOE4 exacerbates synapse loss and neurodegeneration in Alzheimer’s disease patient iPSC-derived cerebral organoids. Nat Commun 11, 5540 (2020).

83. Ashburner, M. et al. Gene Ontology: tool for the unification of biology. Nat Genet 25, 25–29 (2000).

84. Da Cruz, S. & Cleveland, D. W. Understanding the role of TDP-43 and FUS/TLS in ALS and beyond. Curr Opin Neurobiol 21, 904–919 (2011).

85. Vaquero-Garcia, J. et al. RNA splicing analysis using heterogeneous and large RNA-seq datasets. Nat Commun 14, 1230 (2023).

86. Zhang, D. et al. Neuregulin-3 (NRG3): A novel neural tissue-enriched protein that binds and activates ErbB4. Proc Natl Acad Sci U S A 94, 9562–9567 (1997).

87. Buonanno, A. & Fischbach, G. D. Neuregulin and ErbB receptor signaling pathways in the nervous system. Current Opinion in Neurobiology 11, 287–296 (2001).

88. Carteron, C., Ferrer-Montiel, A. & Cabedo, H. Characterization of a neural-specific splicing form of the human neuregulin 3 gene involved in oligodendrocyte survival. J Cell Sci 119, 898–909 (2006).

89. Mei, L. & Nave, K.-A. Neuregulin-ERBB signaling in nervous system development and neuropsychiatric diseases. Neuron 83, 27–49 (2014).

90. Ou, G., Lin, W. & Zhao, W. Neuregulins in Neurodegenerative Diseases. Front. Aging Neurosci. 13, (2021).

91. Takahashi, Y. et al. ERBB4 Mutations that Disrupt the Neuregulin-ErbB4 Pathway Cause Amyotrophic Lateral Sclerosis Type 19. Am J Hum Genet 93, 900–905 (2013).

92. Kao, W.-T. et al. Common genetic variation in Neuregulin 3 (NRG3) influences risk for schizophrenia and impacts NRG3 expression in human brain. Proceedings of the National Academy of Sciences 107, 15619–15624 (2010).

93. Exposito-Alonso, D. et al. Subcellular sorting of neuregulins controls the assembly of excitatory-inhibitory cortical circuits. eLife 9, e57000 (2020).

94. Zhang, Y. et al. Rapid Single-Step Induction of Functional Neurons from Human Pluripotent Stem Cells. Neuron 78, 785–798 (2013).

95. Sholl, D. A. Dendritic organization in the neurons of the visual and motor cortices of the cat. J Anat 87, 387–406.1 (1953).

96. Ferreira, T. A. et al. Neuronal morphometry directly from bitmap images. Nat Methods 11, 982–984 (2014).

97. Wu, Y., Wang, J. & Zhao, Q. Advancements in TDP-43 research: Towards biomarkers and therapeutic targets for amyotrophic lateral sclerosis. Aging and Health Research 5, 100215 (2025).

98. Lukavsky, P. J. et al. Molecular basis of UG-rich RNA recognition by the human splicing factor TDP-43. Nat Struct Mol Biol 20, 1443–1449 (2013).

99. Shiura, H. et al. PEG10 viral aspartic protease domain is essential for the maintenance of fetal capillary structure in the mouse placenta. Development 148, dev199564 (2021).

100. Ono, R. et al. Deletion of Peg10, an imprinted gene acquired from a retrotransposon, causes early embryonic lethality. Nat Genet 38, 101–106 (2006).

101. Shiura, H. et al. PEG10-ORF1 programs trophoblast progenitor development for placental labyrinth formation. 2025.10.08.681076 Preprint at 10.1101/2025.10.08.681076 (2025).

102. Kim, S. H. et al. Axon guidance genes modulate neurotoxicity of ALS-associated UBQLN2. eLife 12, e84382 (2023).

103. Gorrie, G. H. et al. Dendritic spinopathy in transgenic mice expressing ALS/dementia-linked mutant UBQLN2. Proceedings of the National Academy of Sciences 111, 14524–14529 (2014).

104. Lin, B. C., Higgins, N. R., Phung, T. H. & Monteiro, M. J. UBQLN proteins in health and disease with a focus on UBQLN2 in ALS/FTD. The FEBS Journal 289, 6132–6153 (2022).

105. Zheng, T., Yang, Y. & Castañeda, C. A. Structure, dynamics and functions of UBQLNs: at the crossroads of protein quality control machinery. Biochem J 477, 3471–3497 (2020).

106. Davis, M. W. & Jorgensen, E. M. ApE, A Plasmid Editor: A Freely Available DNA Manipulation and Visualization Program. Front Bioinform 2, 818619 (2022).

107. Schindelin, J. et al. Fiji: an open-source platform for biological-image analysis. Nat Methods 9, 676–682 (2012).

108. Bolger, A. M., Lohse, M. & Usadel, B. Trimmomatic: a flexible trimmer for Illumina sequence data. Bioinformatics 30, 2114–2120 (2014).

109. Dobin, A. et al. STAR: ultrafast universal RNA-seq aligner. Bioinformatics 29, 15–21 (2013).

110. Liao, Y., Smyth, G. K. & Shi, W. The Subread aligner: fast, accurate and scalable read mapping by seed-and-vote. Nucleic Acids Res 41, e108 (2013).

111. R Core Team. R: A Language and Environment for Statistical Computing. R Foundation for Statistical Computing. https://doi.org/10.32614/R.manuals (2026) doi:10.32614/R.manuals.

112. Love, M. I., Huber, W. & Anders, S. Moderated estimation of fold change and dispersion for RNA-seq data with DESeq2. Genome Biol 15, 550 (2014).

113. Vaquero-Garcia, J. et al. A new view of transcriptome complexity and regulation through the lens of local splicing variations. eLife 5, e11752 (2016).

114. Robinson, J. T. et al. Integrative Genomics Viewer. Nat Biotechnol 29, 24–26 (2011).

115. Thorvaldsdóttir, H., Robinson, J. T. & Mesirov, J. P. Integrative Genomics Viewer (IGV): high-performance genomics data visualization and exploration. Brief Bioinform 14, 178–192 (2013).

